# Machine Learning Identifies Signatures of Macrophage Reactivity and Tolerance that Predict Disease Outcomes

**DOI:** 10.1101/2022.06.27.497783

**Authors:** Pradipta Ghosh, Saptarshi Sinha, Gajanan D. Katkar, Daniella Vo, Sahar Taheri, Dharanidhar Dang, Soumita Das, Debashis Sahoo

## Abstract

Single-cell transcriptomic studies have greatly improved organ-specific insights into macrophage polarization states are essential for the initiation and resolution of inflammation in all tissues; however, such insights are yet to translate into therapies that can predictably alter macrophage fate. Using machine learning algorithms on human macrophages, here we reveal the continuum of polarization states that is shared across diverse contexts. A path, comprised of 338 genes accurately identified both physiologic and pathologic spectra of “*reactivity*” and “*tolerance*”, and remained relevant across tissues, organs, species and immune cells (> 12,500 diverse datasets). This 338-gene signature identified macrophage polarization states at single-cell resolution, in physiology and across diverse human diseases, and in murine pre-clinical disease models. The signature consistently outperformed conventional signatures in the degree of transcriptome-proteome overlap, and in detecting disease states; it also prognosticated outcomes across diverse acute and chronic diseases, e.g., sepsis, liver fibrosis, aging and cancers. Crowd-sourced genetic and pharmacologic studies confirmed that model-rationalized interventions trigger predictable macrophage fates. These findings provide a formal and universally relevant definition of macrophage states and a predictive framework (http://hegemon.ucsd.edu/SMaRT) for the scientific community to develop macrophage-targeted precision diagnostics and therapeutics.

**One Sentence Summary:** *<u>S</u>*ignatures of *<u>ma</u>*crophage *r*eactivity and *t*olerance (*SMaRT*) predict disease outcomes

## INTRODUCTION

Macrophages are complex; as sentinel cells of the innate immune system, they are found in various organs and their dysregulated activation can directly impact organ functions and the outcome of all diseases ^1,2^. Macrophages were initially classified as M1 (the classically activated macrophages) and M2 (the alternatively activated macrophages) based on their functions at the extremes of polarization states ^3^. However, the M1/M2 nomenclature is considered as too simplistic; it fails to describe the diverse, polyfunctional and plastic cells, and the myriad of continuum states that they adopt in the tissue at steady-state and during disease ^4-7^. To cope with this degree of diversity and plasticity, several definitions of macrophage subtypes have emerged, each representing specialized contexts, e.g., TAMs, tumor-associated macrophages ^8^; LAMs, lipid-associated macrophages in atherosclerosis ^9^; DAMs, disease-associated microglia in neurodegenerative disorders ^10^; SAMs, scar-associated macrophages in liver fibrosis ^11-13^. These definitions were geared to identify divergent markers, spatial localization, origin, and functional pathways associated with macrophages during disease; however, they fall short in predictive or prognostic abilities.

We sought to create and validate a comprehensive model of macrophage processes for defining, tracking, and even predicting macrophage fate after perturbation (see **Fig 1a, S1A** for workflow outline). We hypothesized that such a model might inspire *formal* definitions for macrophage polarization states that are reflective of fundamental processes and maintain relevance across tissues, organs, diseases and species. In addition, it may also rationalize diagnostics and therapeutics to detect and reset, respectively, deranged macrophage states in disease. We show that such formal definition(s) of macrophage states is not only possible, but also provide evidence for their usefulness in single cell data analysis, in prediction and prognostication.

**Figure 1:**
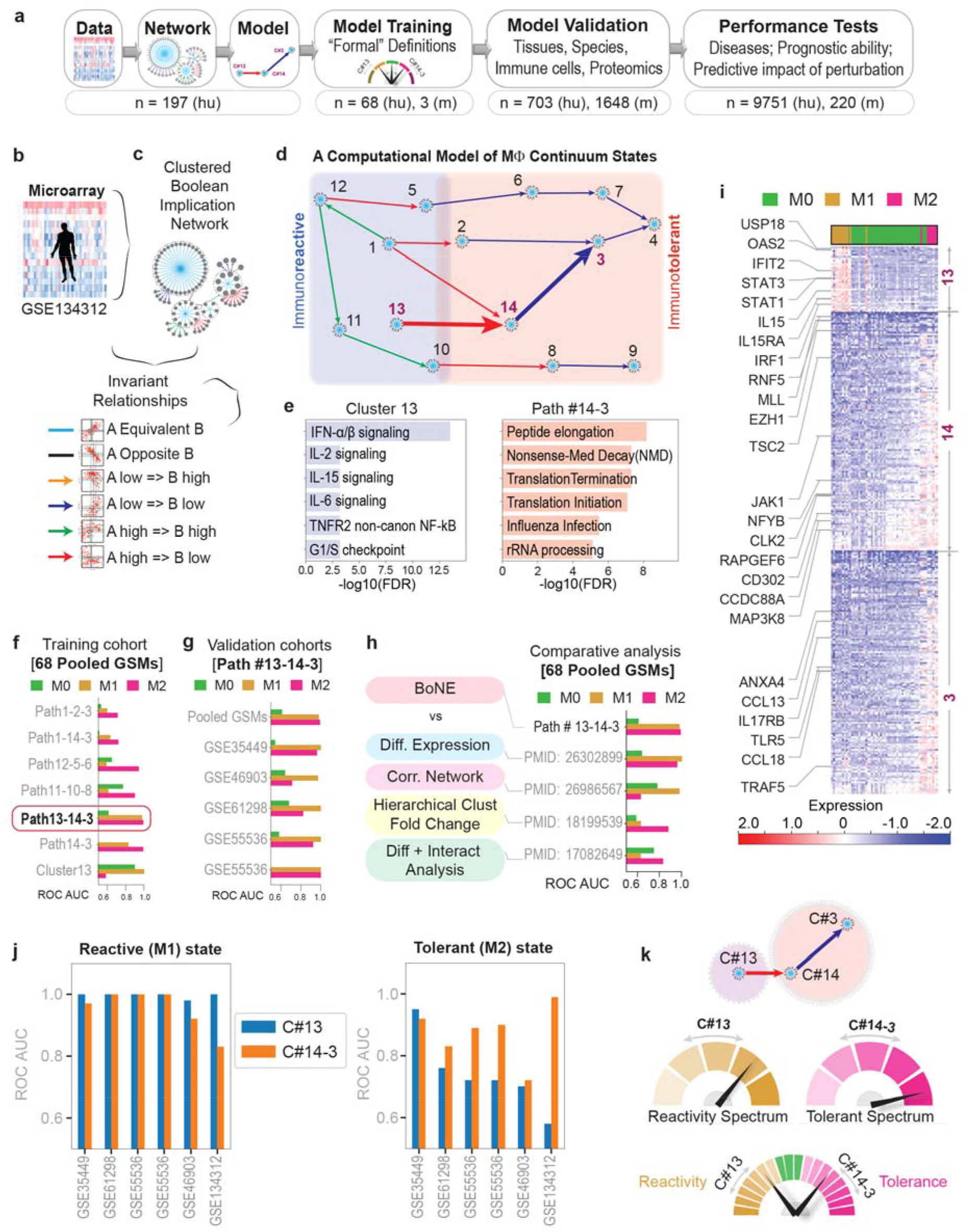
*BoNE*-assisted formulation of formal definitions of macrophage polarization. **a)** Overview of workflow and approach used in this work. **b-c)** A pooled dataset of diverse human transcriptomes (**b**; n = 197) was used to build a Boolean implication network (**c**-*top*) and visualized as gene clusters (nodes, comprised of genes that are equivalent to each other) that are interconnected based on one of the six overwhelming Boolean implication relationship between the clusters (directed edges; **c**-*bottom*). **d)** Display of the major Boolean paths within the network prioritized based on the cluster size. Annotations of “immunoreactive” and “immunotolerant” ends of the spectrum are based on the expression profile of the gene clusters in 68 samples within the pooled dataset that were stimulated *in vitro* as M1 and M2, respectively. **e)** Reactome pathway analysis of each cluster along the top continuum paths was performed to identify the enriched pathways (for other clusters see http://hegemon.ucsd.edu/SMaRT/). **f-g)** Training (**f**) was performed on the 68 pooled samples using machine-learning approaches; the best-performing Boolean path, #13-14-3 was then validated (**g**) in multiple independent human macrophage datasets. For a list of datasets used see **Table S1**. The performance was measured by computing *ROC AUC* for a logistic regression model. **h)** Comparative analysis of performance of the *BoNE*-derived *versus* other traditional approaches in segregating M0/M1/M2 polarization states. **i)** Heatmap displaying the pattern of gene expression in C#13, 14 and 3. Selective genes are labelled. **j)** Validation studies assessing the ability of the genes in either C#13 alone or C#14-3 alone to classify M0/M1/M2 polarization states in multiple human macrophage datasets. **k)** *<u>Top</u>*: Schematic summarizing the model-derived formal definitions of macrophage polarization states based on the levels of expression of genes in C#13 (hypo to hyper-“reactivity” spectrum) and those in C#14+3 (hypo to hyper-“tolerant” spectrum). *<u>Bottom</u>*: A composite score of the entire range of physiologic and pathologic response can be assessed via the *BoNE*-derived path #13→14→3.

## RESULTS AND DISCUSSION

### A computational model of continuum states in macrophage processes

We chose a Boolean approach to build transcriptomic network ^14^; this approach has been used to create maps of evolving cellular states along any disease continuum and identify cellular states in diverse tissues and contexts with high degrees of precision (see detailed *Methods*). The Boolean approach relies on invariant relationships that are conserved despite heterogeneity in the samples used for the analysis. Invariant relationships among pairs of genes that are conserved across samples representative of maximum possible diversity, i.e., irrespective of their origin (normal or disease), laboratories/cohorts, different perturbations, are assumed to be fundamentally important for any given process.

For model training and development, we used a pooled all-human microarray dataset that included 197 manually annotated heterogeneous macrophage datasets from GEO (<u>GSE134312</u> ^15^; **Fig 1a-c**; **Fig S1A-B;** see **Supplemental Information 1** for catalog of datasets). These datasets contained primary tissue-derived macrophages (both healthy and diseased tissues) and cultured macrophage cell lines (e.g., THP1), either untreated or treated with diverse sets of ligands that are known to induce either M1 (n = 13) or M2 (n = 8) polarized states (see **Table S1**).

A graph (**Fig 1d**; **Fig S1C; Fig S2A**) is built, comprised of gene clusters (nodes) connected to each other using Boolean implication relationships (edges). The network displayed scale-free properties, as expected (**Fig S1D**). We oriented ourselves to the resultant network by querying and locating the known ‘M1/M2’ samples; the ‘M1’ samples segregated towards one end, and ‘M2’ samples on the other, implying that the paths of connected clusters within the resultant network represent a continuum of cellular states in macrophages within the immunologic spectrum (**Fig S1E-G**). Reactome pathway analyses ^16^ of each cluster along the top continuum paths revealed a multitude of cellular processes that are impacted during macrophage polarization (**Fig 1e;** Gene clusters and reactome pathways can be queried at: http://hegemon.ucsd.edu/SMaRT/).

### Identification of <u>s</u>ignatures of <u>ma</u>crophage ‘reactivity’ and ‘tolerance’ (SMaRT)

Using machine learning approaches, various interconnected gene clusters (i.e., Boolean paths) were assessed for their ability to accurately classify the samples (based on the genes in the clusters and computing a weighted average of gene expression values outlined in **Fig S2B**) (**Fig 1f**). A multivariate analysis of the top five Boolean paths revealed that the path connecting clusters(C)#13→14→3 is the best (p < 0.001) at discriminating M1 (ROC-AUC 0.98) and M2 (ROC-AUC 0.99) (**Fig 1f; Fig S2C**). Path #13→14→3 was subsequently validated in five other independent datasets (**Fig 1g**). A comparative analysis of #13→14→3 path *vs*. other traditional approaches, e.g., Differential Expression ^17^, Correlation Network ^17^, Hierarchical Clustering ^18^ and Differential and interactome analyzes ^19^ showed the superiority of the *BoNE*-derived path in separating M0-M1-M2 states. The Boolean path matched differential expression in its ability to distinguish M1 state, while exceeding the remaining traditional approaches (**Fig 1h**). A heatmap of the pattern of gene expression in each cluster in M0-M1-M2 states is shown in **Fig 1i**.

Furthermore, C#13 predicted M1 perfectly (ROC-AUC = 1.0); the path #14→3 predicted M2 close to perfection (ROC-AUC = ranging from 0.80 to 1.00) in all cohorts tested (**Fig 1j**). This indicates that while the path #13→14→3 is the most accurate path across all human macrophage-derived datasets collected and analyzed, C#13 and the path #14→3 carry relevant information on macrophage states independently of each other. C#13 is associated with M1-like state and expression of these genes is predicted to reflect the extent of “immunoreactivity” of macrophages. Path #14→3 is associated with a M2-like state and expression of these genes is predicted to reflect the extent of “immunotolerance”. We define the two distinct macrophage polarization states in physiology as *“*reactive” and “tolerant” based on basal C#13 and #14→3 scores, respectively (**Fig. 1k**). Four additional macrophage states could also exist, presumably in disease states, i.e., hyperreactive (high C#13), hypertolerant (high #14→3), hyporeactive (low C#13), and hypotolerant (low #14→3) (**Fig. 1k**). Henceforth, we refer to these genes as *<u>s</u>*ignatures of *<u>ma</u>*crophage *r*eactivity and *t*olerance, abbreviated as ‘*SMaRT’* (See http://hegemon.ucsd.edu/SMaRT/ and **Supplemental Information 2** for the list of genes).

### SMaRT genes are relevant across tissues, organs, species and immune cells

We found that the path #13→14→3 successfully identified M1/M2-polarization states in diverse tissue-resident macrophages (brain-resident microglia, the Langerhan’s cells in the skin, intestinal and lung alveolar macrophages, etc.), in both humans and mice (**Fig 2a-b**). See **Supplemental Information 1** for the degree of heterogeneity represented in these datasets. Surprisingly, the path could also separate reactive and tolerant states of other immune cells, including lymphocytes (B/T and NK-T), natural killer (NK) cells, neutrophils, dendritic, basophils, eosinophils and mast cells (**Fig 2c**). Together, these findings indicate that the *SMaRT*-based definitions of ‘reactivity’ and ‘tolerance’ remain relevant in the context of tissue-resident macrophages despite their adaptation to the tissue and/or organ-specific microenvironment for their identity ^20-22^. These definitions also maintain relevance in mice, whose immune system is different from ours ^23^. Findings suggest that the *SMaRT*-based definitions may reflect the *fundamental* immune-reactive and tolerant gene regulatory mechanisms that are shared among diverse cells in our immune system, regardless of whether they are derived from the myeloid or lymphoid lineage (**Fig 2d**).

**Figure 2.**
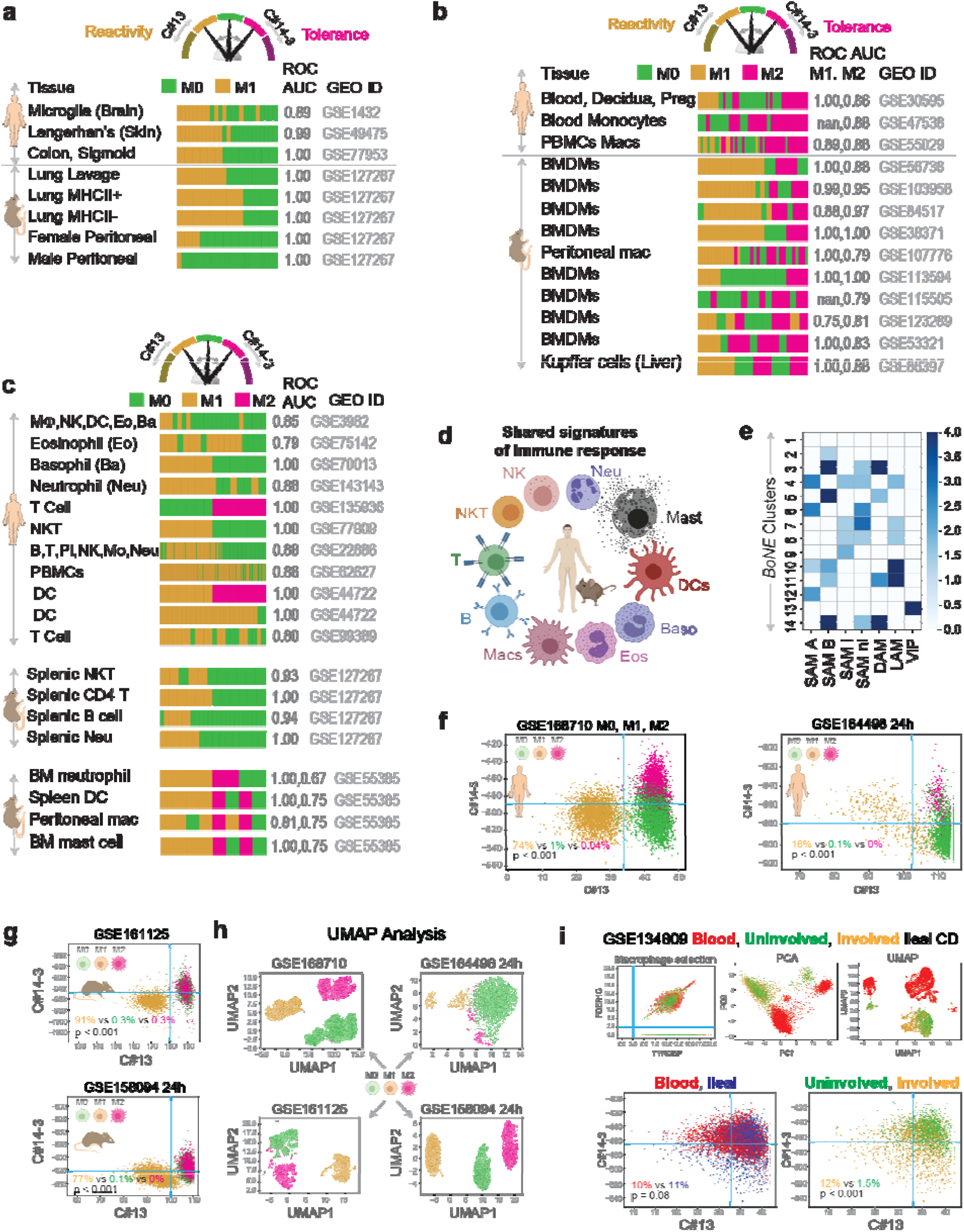
Definitions of “reactivity” and “tolerance” are conserved across tissues, organs, species and diverse immune cell types. **a-b**) Validation studies assessing the ability of *SMaRT* genes to classify diverse tissue-resident macrophage datasets from both humans and mice. Performance is measured by computing *ROC-AUC*. Barplots show the ranking order of different sample types based on the composite scores of C#13 and path #14-3. **c-d**) Validation studies (c) assessing the ability of *SMaRT* genes to classify active vs inactive states of diverse immune cell types in both humans and mice. The schematic (d) summarizes findings in c. **e**) Published disease-associated macrophage gene signatures (see **Supplemental Information 2**) are are analyzed for significant overlaps with various gene clusters in the Boolean map of macrophage processes. Results are displayed as heatmaps of -Log10(*p)* values as determined by a hypergeometric test. **f-g**) Scatterplots of the composite score of C#13 and path #14-3 in human (f, GSE168710, GSE164498 24h) and mouse (g, GSE161125, GSE158094 24h) single cell RNASeq datasets with well defined macrophage polarization states (M0, M1, M2). Blue lines corresponds to the StepMiner thresholds. Percentages of different cell types are reported in the bottom-left quadrant. Pvalue is computed by two tailed two proportions z-test for M1 vs M0. **h)** Traditional UMAP analysis of the single cell RNASeq datasets. **i)** PCA, UMAP and BoNE analysis of single cell RNASeq dataset GSE134809 that includes blood and ileal biopsy (uninvolved and involved) samples from Crohn’s disease (CD) patients. Macrophages were selected as the top right corner by using thresholds (2.5, blue lines) on TYROBP and FCER1G. Blue lines correspond to the StepMiner thresholds in the scatterplot between C#13 and C#14-3 (bottom plots). Bottom-left quadrant is evaluated for enrichment of cell types across tissue (blood vs ileal) and disease states (uninvolved vs involved CD). Percentages of different cell types are reported in the bottom-left quadrant. P value is computed by two tailed two proportions z-test.

### The network captures physiologic macrophage states and functions

We found that diverse macrophage subtypes are represented within our model of macrophage processes (**Fig S2I**). The classical M1 subtype was represented in C#1 and #13 on the reactive end of the model, alongside TCR+ macrophages in C#1 and #12; the latter is known to release CCL2 and have high phagocytic abilities ^24^. On the tolerant end of the model we found the TAMs in C#2, #5, #6 and the *CD169*+ macrophages in C#2, #3 and #7; both subtypes have been implicated in immunological tolerance ^25-27^. As one would anticipate, the tissue-resident macrophages (M2a-d) that are known for their plasticity of polarization states were more centrally placed in C#2 and #5. Finally, gene signatures of scar-associated non-inflammatory (ni) macrophages that restrict inflammation in liver cirrhosis (SAM B ^12^ and SAM ni ^13^, **Fig 2e**) and damage-associated microglia (DAMs ^10^; **Fig 2e**) that restrict the progression of neurodegeneration significantly overlapped with the tolerant clusters C#14 and #3. A gene signature that was recently shown to be induced in monocytes and macrophages in all viral pandemics (ViP), past and present, overlapped with the reactive C#13 as expected (**Supplemental Information 2** lists all gene signatures in **Fig 2e**).

Members of the family of pattern recognition receptors (PRRs; **Table S2**), *via* which macrophages ‘sense’ its surroundings ^28^, were distributed in various nodes within the model, overlapping with each other (**Fig S2J**). PRRs that sense pathogens or apoptotic cells to stimulate phagocytosis and mediate inflammation, e.g., toll-like (TLRs), nucleotide oligomerization domain (NODs) and receptor for advanced glycation end products (RAGE) were found on the ‘reactive’ side of the model. The TLRs, scavengers and C-type lectins also overlapped with path#13→14→3, but only on the tolerant end (cluster #3) of the spectrum.

The circadian genes were distributed within clusters along a path (#1→2→3→4) (**Fig S2K**), intersecting at the tolerant end of the path#13→14→3, i.e., C#3. The daytime circadian genes were in the reactive end of the model and showed an inverse high=>low Boolean relationship with night-time circadian genes; the latter were mostly in the tolerant end of the model (**Fig S3A-C**). This finding is consistent with the current belief that macrophages ‘kill’ (reactive) during the day and ‘heal’ (tolerant) during the night ^29^. We also show that the performance of the tolerant signature (C#14-3) in diseases that have an intricate relationship with circadian rhythms, such as metabolic syndrome ^30^, can be further improved by normalization based on a clock gene or clock gene signature (**Fig S4**).

### SMaRT genes identify polarization states at single cell resolution

Analysis of C#13 vs path #14→3 composite scores revealed a consistent pattern of macrophages polarized towards M1 and M2 in multiple independent single cell RNASeq data (**Fig 2f-g, Fig S2D-G**) in both human (<u>GSE168710</u>, <u>GSE164498</u>, **Fig 2f**) and mouse (<u>GSE161125</u>, <u>GSE158094</u>, **Fig 2g**). StepMiner threshold (blue lines, **Fig 2f-g**) is computed for both C#13 and path #14→3 composite scores that divide the scatterplots into four different quadrants. Bottom-left quadrant shows a significant enrichment of reactive macrophages (M1) in all four scatterplots (p < 0.001, **Fig 2f-g**). Traditional UMAP analyses in the above datasets show distinct clusters for the different polarized states (**Fig 2h**) but it is hard to translate that in Crohn’s disease (CD) dataset (<u>GSE134809</u>, **Fig 2i***-top*). Since BoNE derived signatures show significant enrichment of the reactive macrophages in the bottom-left quadrant (as shown in **Fig 2f-g**), it can easily be tested in CD dataset. As expected, in the bottom-left quadrant (reactive macrophages) of the CD dataset (**Fig 2i***-bottom*), macrophages from involved tissues are significantly enriched (p < 0.001) compared to uninvolved where as no significant enrichment (p = 0.08) was observed between ileal vs blood tissue. These findings are consistent with macrophage phenotypes observed in inflammatory bowel disease patients ^31,32^.

Furthermore, to test if presence of both reactive and tolerant states can be detected by BoNE models, we artificially created pseudobulk samples using various proportions of of M1, M2 cells, and one sample with 30% M1 and 30% M2 cells (Mixed) in the background of lung cells from a single cell RNASeq dataset <u>GSE150708</u> (human, **Fig S2D**). The Mixed sample was categorized as both tolerant and reactive as expected using C#13 and path #14-3 signatures (**Fig S2D**). The blood tissue from CD dataset (<u>GSE34809</u>, **Fig S2G**) is categorized as both tolerant and reactive which may be interpreted as presence of these mixed states.

### SMaRT genes identify pathologic polarization states in diseases

We next asked how the Boolean network-derived *formal* definitions perform in disease states. A plethora of disease conditions and tissues were analyzed (**Fig 3a-n; Supplemental Information 1**). We computed a composite immune response score derived from C#13 alone or C#14 and #3, which quantitatively estimates the degree of “reactivity” and “tolerance”, respectively, and tested it in diverse conditions. An analysis of full-thickness colon tissues representing the 2 major subtypes of inflammatory bowel disease (IBD), ulcerative colitis (UC) and Crohn’s disease (CD) (**Fig S5A**) revealed that reactivity is a common feature in both UC and CD (**Fig 3a**, *top*; **Fig S5B**-*left*). However, tolerance was enhanced only in CD (**Fig S5B***-right*), which is consistent with the notion that ‘alternatively’ activated tolerant macrophages may drive the transmural nature of the inflammation, ineffective bacterial clearance, and accompanying tissue remodeling (fibrosis, stricture, fistula), all features that are observed uniquely in CD ^33^, but not UC. Reactivity alone could prognosticate outcome (i.e., segregate responder *vs* non-responder) regardless of the heterogeneity of the UC cohorts and the diverse treatment modalities (**Fig S5C-D**), consistent with the widely-accepted notion that hyperinflammatory macrophages are drivers ^34^ of the disease and key targets for therapeutics ^35^. Insufficient datasets precluded similar analyses in the case of CD.

**Figure 3.**
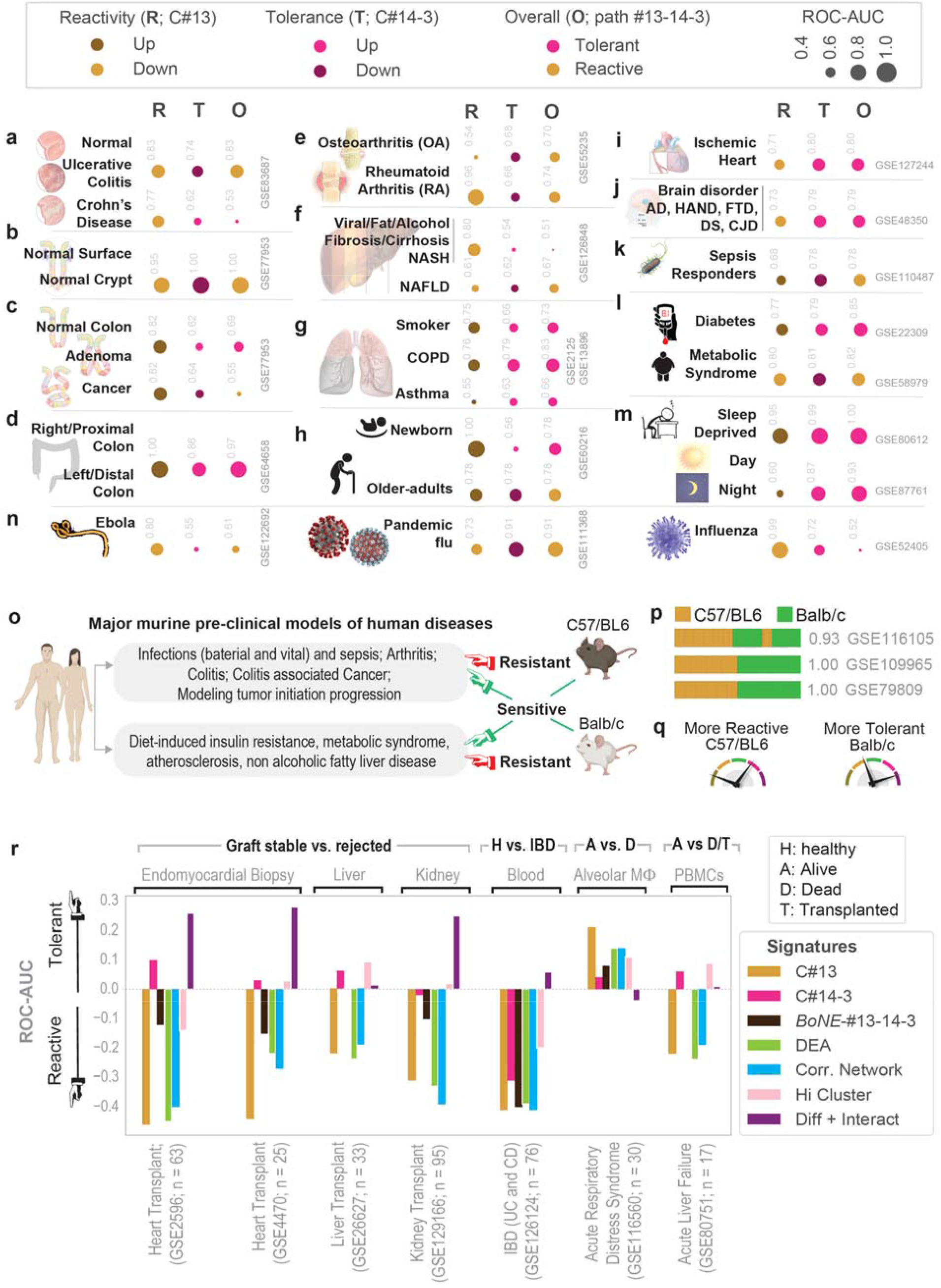
Definitions of “reactivity” and “tolerance” detects pathologic macrophage states in disease. Tissue immune microenvironment is visualized (in panels a-n) as bubble plots of ROC-AUC values (radii of circles are based on the ROC-AUC; Key on top) demonstrating the direction of gene regulation (Up vs Down; Key on top) for the classification of samples using BoNE-derived gene signatures of either reactive (R; C#13) or tolerant (T; C#14-3) or overall (O; path #13→14→3) in columns. The ROC-AUC values are provided next to the bubble. Sample diversity and sizes are as follows: **a)** IBD; GSE83687, n = 134; 60 Normal, 32 Ulcerative Colitis, 42 Crohn’s Disease. **b)** Colon crypt; GSE77953, 6 Normal Surface *vs*. 7 Normal Crypt base. **c**; Colon cancer: Pooled colon dataset from NCBI GEO; n=170 Normal, 68 Adenomas, 1662 CRCs. **d)** Colon anatomy: Proximal (right) vs distal (left) normal colon from mouse (GSE64423, n = 6) and human (GSE20881, n = 75). See **Figure S5** for violin plots. **e)** Arthritis; GSE55235, GSE55457 and GSE55584, n = 79; 20 Normal, 33 Rheumatoid Arthritis, 26 Osteoarthritis. **f)** Hepatitis: GSE89632, n=63; 20 fatty liver, 19 Non-alcoholic steatohepatitis (NASH) and 24 healthy, alcoholic liver disease (GSE94417, GSE94397 and GSE94399, n = 195; 109 Healthy, 13 Alcoholic Hepatitis, 6 Alcoholic fatty liver (AFL), 67 Alcoholic cirrhosis (AC) and viral hepatitis (GSE70779, n=18; 9 Pre-treatment, 9 Post-treatment with direct-acting anti-virals). **g)** Chronic lung disease; GSE2125 and GSE13896, n = 115; 39 Non-smoker, 49 Smoker, 15 Asthma, 12, Chronic Obstructive Pulmonary Disease (COPD). **h)** Aging process; GSE60216, n = 9; 3 Newborn babies, 3 Adults, 3 Old-adults. **i)** Cardiomyopathy (CM), ischemic and non-ischemic (I/NI); GSE104423, n = 25 human samples; 14 NICM, 11 ICM; GSE127244, n = 24 mouse samples, 16 NICM, 8 ICM. **j)** Neurodegenerative brain disorders; GSE118553 (n = 401) and GSE48350 (n = 253), Alzheimer’s disease (AD); GSE35864, HIV-associated neurocognitive disorder (HAND; n = 72); GSE13162, frontotemporal dementia (FTD; n = 56); GSE59630, Down’s Syndrome (DS; n = 116); GSE124571, Creutzfeldt-Jakob Disease (CJD; n = 21). **k)** Systemic inflammatory response syndrome (SIRS) and sepsis; GSE63042 (n = 129); GSE110487 (n = 31). **l)** Type 2 diabetes and metabolic syndrome; GSE22309 (n = 110), Pre- and post-insulin treatment muscle biopsies from 20 insulin sensitive, 20 insulin resistant, 15 T2DM; GSE98895 (n = 40), PBMCs from 20 control, 20 metabolic syndrome. **m)** Sleep deprivation and circadian rhythm; GSE9444, n = 131 mouse brain and liver samples; GSE80612, twin, n = 22 human peripheral blood leukocytes; GSE98582, n = 555 human blood samples; GSE104674, n = 48, 24 healthy and 24 T2DM. **n)** Viral pandemics, such as SARS, MERS, Ebola and others [**N**; numerous datasets, see **Figure S7E**]. See **Figure S6** and **S7** for violin plots relevant to panel **e-n**. **o-q**) Schematic (o) summarizes the use of two major mouse strains (C57/B6 and Balb/c) commonly used for modeling two broad categories of human diseases. Bar plots (p) showing sample classification of genetically diverse macrophage datasets based on expression levels of genes in C#13. Schematic (q) summarizes findings. **r**) The diagnostic potential of various indicated gene signatures were tested on multiple datasets generated from tissues derived from patients with the known clinically relevant outcome, as indicated. In each case, BoNE-derived signatures were compared against four traditional approaches.

We also found that “reactivity” and “tolerance” differs along the length of the colon crypt—the surface is more reactive, whereas the stem-cell niche at the bottom is more “tolerant” (**Fig 3b**; **Fig S5E-F**). We also found that “hypo-reactivity” [low C#13] and “complete tolerance” [high #14→3] are two states that are progressively accentuated during colorectal carcinoma (CRC) initiation and the emergence of chemoresistance (**3c**; **Fig S5G-H**). Consistent with the fact that most of the CRCs are found located in the left (distal) colon and microbe-driven risk is high in that segment ^36^, we found that segment to be more tolerant than the right (proximal) segment (**3d**).

We detected altered macrophage states during the initiation and progression of several human other diseases, ranging from arthritis, through neurodegenerative diseases to viral pandemics (see **Fig 3e-n; Fig S6A-N, Fig S7A-E**). Our definitions for “reactivity” and “tolerance” could accurately identify the underlying pathologic macrophage states implicated in each condition. Together, these results show that the *BoNE*-derived signature can detect different subsets of macrophages are essential to the pathogenesis of many diseases. Findings also agree with the notion that disease chronicity is invariably associated with mixed polarization states (whose detection has largely been enabled by scSeq studies) where each state plays an opposing (balanced) role ^2,8-13^.

### SMaRT genes rationalize the choice of mouse models

Although mice are the preferred model species for research ^37^, most agree that their innate immune systems differ ^23^. C57BL/6J and Balb/c mice are two most commonly used mouse strains that differ in their immune responses, giving rise to distinct disease outcomes, which in turn rationalizes their use as pre-clinical models for human diseases (**Fig 3o**). Our signature successfully classified the macrophages from these two strains in three independent cohorts ^38,39^ (**Fig 3p**); C57BL/6 emerged as more reactive and Balb/c as more tolerant (**Fig 3q**). These findings are consistent with the observation that BALB/c mice are more susceptible to a variety of pathogens ^40-42^, and are useful for modeling tumor initiation and progression and for making antibodies. By contrast, C57BL/6 mice are resistant to infections and are the most common strain used for modeling inflammatory diseases, e.g., arthritis, metabolic disorders [NASH, atherosclerosis, etc. ^43-45^]. We conclude that the model-derived definitions for “reactivity” and “tolerance” — (i) capture the contrasting immunophenotypes of these two murine strains previously reported by Mills et al., ^3^ and (ii) rationalize the choice of each strain as preferred models for modeling a unique set of human diseases. Findings also suggest that the model-derived signatures could serve as an objective guide for assessing the appropriateness of any species/strains/sub-strains as pre-clinical models.

### SMaRT genes carry diagnostic value

Next we compared head-to-head the diagnostic and prognostic potential of the newly defined polarization states against four traditional definitions: differential expression analysis ^17^ (DExp), correlation network ^46^ (CorN), hierarchical clustering + fold change ^18^ (HiClu), and differential + interactome analysis ^19^ (Dif+Int). A composite immune response score derived from C#13 alone, which quantitatively estimates the degree of “reactivity” was tested on multiple datasets generated from tissues derived from patients with known clinically relevant diagnoses. A hyper-reactive state was invariably associated with graft rejection in transplanted hearts, livers and kidneys (**Fig 3r**). A ‘hyper-reactive’ state also classified IBD-afflicted children from those with non-IBD indications (8-18 y age) with reasonable accuracy in a prospective study where the blood samples were drawn at the time of diagnostic colonoscopy (**Fig 3r**). Among the critically ill patients in the ICU, a hyper-reactive state was associated with better 28-day survival for those with ARDS on ventilators (**Fig 3r**) and improved survival without the need for liver transplantation in those diagnosed with Tylenol-induced acute liver failure (**Fig 3r**). While some of the four other traditional methodologies fared similar to the new definitions in some cohorts, none performed as well, and/or as consistently. Findings suggest that the *BoNE*-derived signatures may capture fundamental aspects of macrophage polarization that drive disease states.

### SMaRT genes can prognosticate outcome

We next computed a composite immune response score based on either the path #13-14-3 or C#13 alone. When used as a composite score, a low score value represents “reactive” and high score value represent “tolerant” states. This signature was tested on all transcriptomic datasets found on the NCBI GEO database (as of *04/2022*) originating from prospective studies, regardless of disease. Prospective studies were chosen because they rarely have selection bias from enrollment procedures because the *outcomes* have not yet occurred at the time of enrollment. In the context of cancers, “reactive” tumors carried a worse prognosis than “tolerant” ones across a variety of solid tumor subtypes, e.g., colorectal (n = 555; **Fig 4a**), breast, pancreas, prostate, glioblastoma and bladder cancers (**Fig S7F**). These findings are consistent with the well-recognized role of inflammatory cells in the tumor microenvironment ^47^.

**Figure 4.**
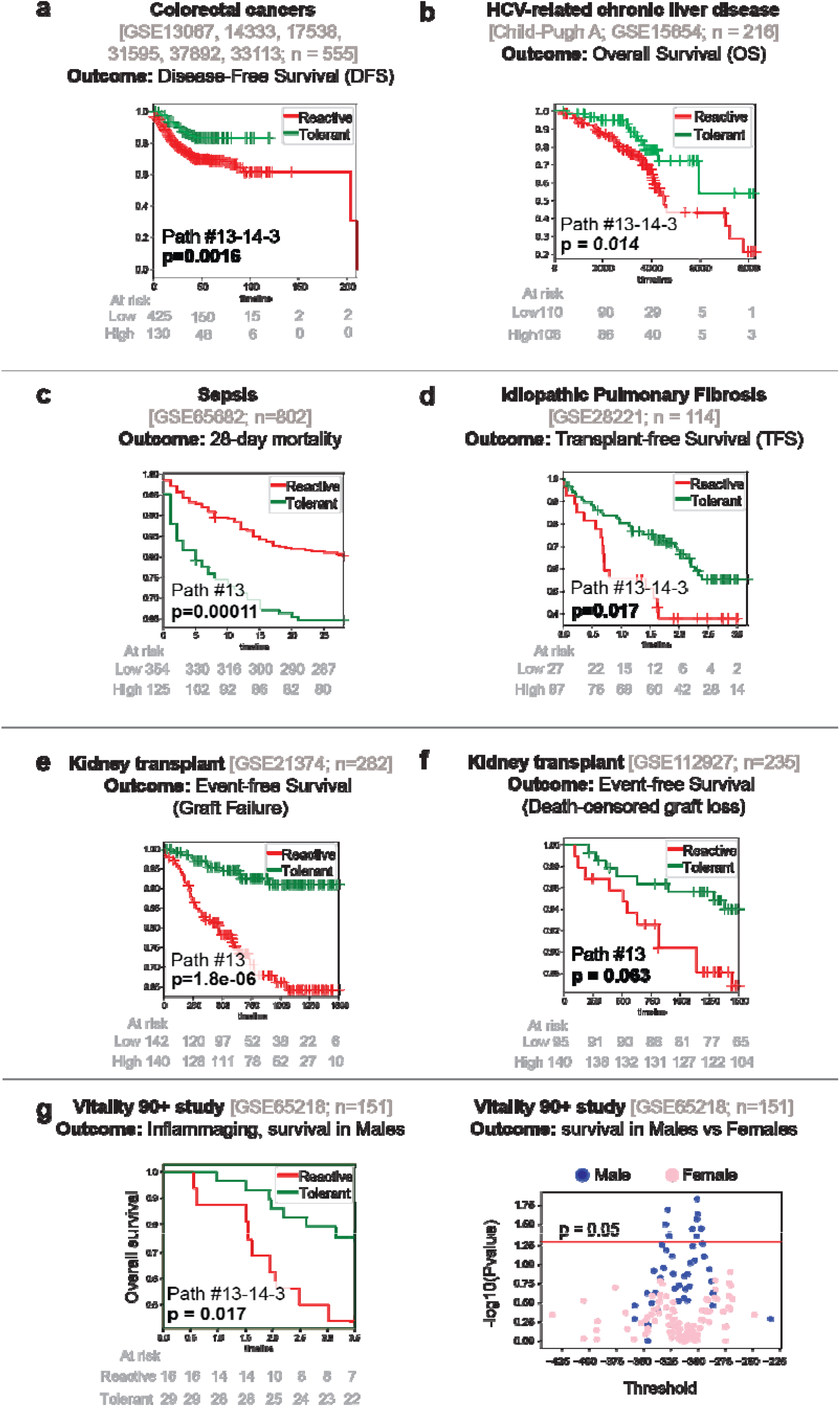
Prognostic potentials of *BoNE model*. **a-g**) The prognostic performance of the *BoNE*-derived *SMaRT* genes is evaluated across diverse disease conditions (colon cancer, a; liver fibrosis, b; sepsis, c; idiopathic pulmonary fibrosis, d; kidney transplantation, e-f; inflammaging, g). Results are displayed as Kaplan Meier (KM) curves with significance (*p* values) as assessed by log-rank-test. A composite immune response score is computed using Boolean path #13→14→3 or C#13 alone, as indicated within each KM plot. Low score = “reactive”; high score = “tolerant”. A threshold is computed using StepMiner by searching three options (thr, thr +/-noise margin) on the immune score to separate these two states. **g-*right***) Scatterplot between all possible thresholds of the #13→14→3 composite score and -log10 of the pvalue from the log-rank test for both male (blue) vs female (pink) separately. Pvalues are significant above the red line (p = 0.05). See also **Fig S7F** for other cancers.

In a cohort of 216 patients with HCV-related liver fibrosis, overall survival was reduced among patients with a “reactive” signature on their liver biopsies compared to those with a “tolerant” signature (**Fig 4b**). These findings are consistent with the known role of activated macrophages in chronic liver injury, inflammation and fibrosis ^48-51^.

In a cohort of 802 patients with sepsis, 28-day mortality was worse among those with a “tolerant” signature compared to those with a “reactive” signature (**Fig 4c**). This finding is consistent with the notion that “endotoxin tolerance” during sepsis carries poor outcome ^52^.

In a cohort of 114 patients with idiopathic pulmonary fibrosis (IPF), an incurable disease that is characterized by progressive fibrosis requiring lung transplantation ^53^, a “reactive” signature was associated with shorter transplant-free survival (**Fig 4d**). Results are in keeping with the widely accepted notion that proinflammatory pulmonary macrophages are known to drive inflammation and fibrosis in the lung ^54^.

Among 517 recipients of kidney transplants, a “reactive” signature was associated with increased graft loss in two independent cohorts (**Fig 4e-f**). Findings are in keeping with prior body of work implicating inflammatory macrophages (both number and extent of activation) as culprits in both acute and chronic allograft rejection and graft loss ^55-57^.

Finally, among 151 nonagenarians in the Vitality 90+ study ^58^, a “reactive” signature was associated with higher mortality in men (**Fig 4g-***left*). No significant results were found in women (**Fig 4g-***right*). Results are in keeping with the fact that the plasma levels of the ‘classical’ marker of inflammaging, i.e., interleukin-6 (IL-6) and a pro-inflammatory gene signature in PBMCs were correlated in men, whereas no correlations were observed in women ^59^.

These findings demonstrate a degree of robustness and consistency in the prognostic ability of the newly defined signatures of macrophage polarization across diverse diseases and independent datasets.

### SMaRT genes are significantly enriched in the macrophage proteome

We used Tandem Mass Tag (TMT) proteomics datasets from THP1-derived macrophages (M0, PMA) that were polarized to M1-M2 states (see workflow **Fig 5a**) and asked if the *BoNE*-derived gene clusters are translated to proteins. We found that the *BoNE*-derived *SMaRT* genes were induced significantly in the THP1 proteome (**Supplemental Information 3**). Consistent with our hypothesis that C#13 and path #14→3 carry independent information regarding “reactivity” and “tolerance”, we found that LPS and IFNγ-induced M1 polarization was associated with significant differential translation of genes in C#13 (**Fig 5b***-top*), whereas IL4-induced polarization was associated with significant differential translation of genes in C#14 and C#3 (**Fig 5b***-bottom*). Such differential protein translation continued to take place over 24 h (**Fig 5b**).

**Figure 5.**
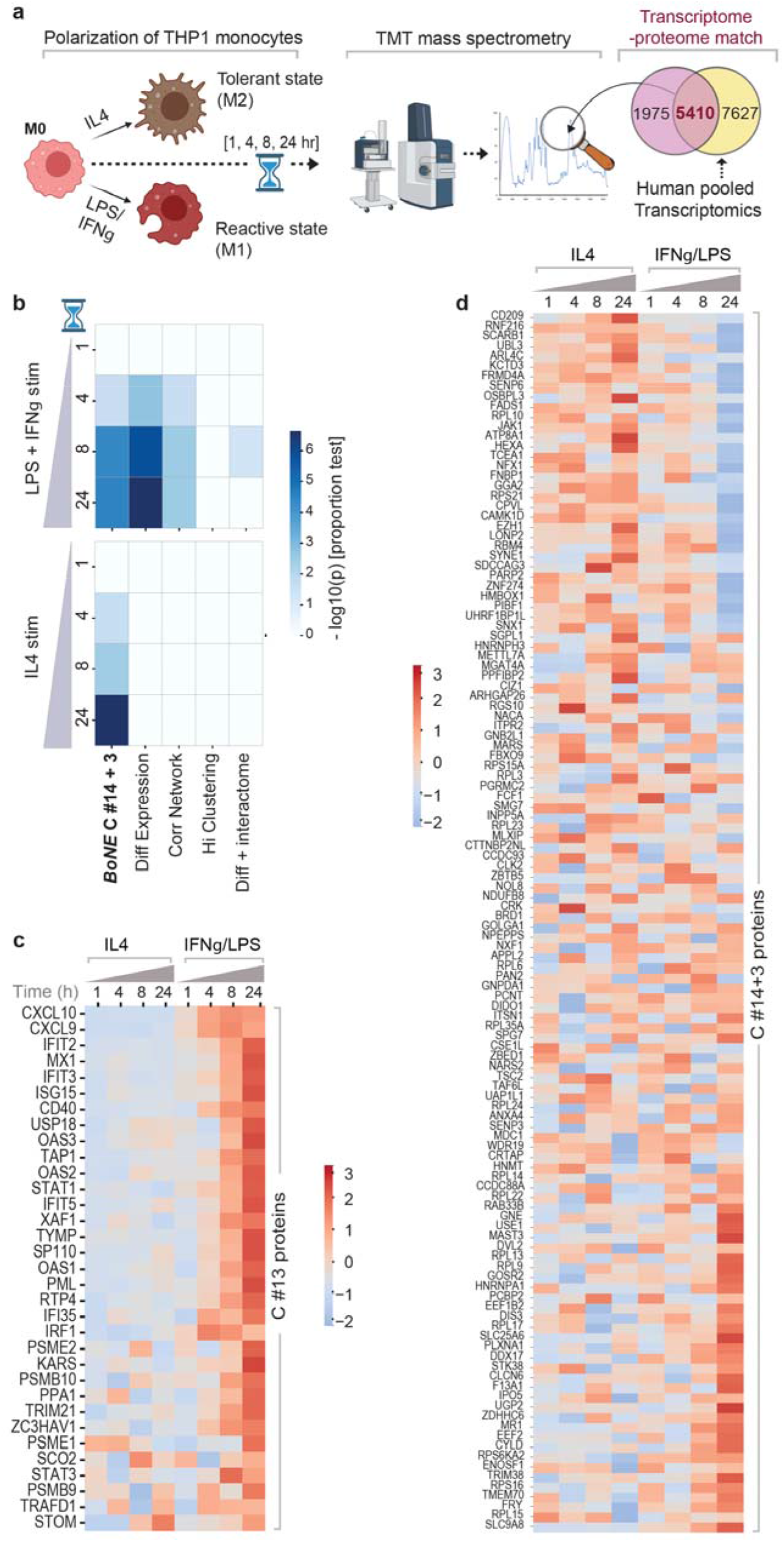
*SMaRT* genes are differentially translated in polarized macrophages. **a)** Overview of the experimental design. PMA-treated human THP-1 cell lines (M0) are polarized to M1 (with LPS and IGNg) or M2 (with IL4), followed by multiplexed mass spectrometry at indicated time points. The fraction of the global macrophage transcriptome (from the pooled 197 macrophage datasets) that is represented in the global macrophage proteome is subsequently assessed for induction (or not) of proteins that are translated by various gene signatures. **b)** Selectivity of induction of proteins upon LPS and IFNg (top) or IL4 (bottom) stimulation at various timepoints was assessed across different signatures using z-test of proportions and -log(10)p values are displayed as heatmaps. **c-d**) z normalized Log of intensities of proteins (Supplemental Information 3) translated at different time points by genes in C#13 (c) and C#14+3 (d) are displayed as heatmaps.

Comparative analyses showed that while the “reactivity” signatures identified by two other conventional methodologies--Differential Expression and Correlation Network--also reached significance; **Fig 5b***-top*), “tolerance” signatures derived by all other conventional approaches did not (**Fig 5b***-bottom*). Heatmaps show the dynamic and opposing nature of the proteins translated by the genes within the *BoNE*-derived gene signatures during polarization (**Fig 5c-d**).

Findings demonstrate that the gene signatures of ‘reactivity’ and ‘tolerance’ identified here are significantly represented also in the translated proteome.

### Perturbation of SMaRT genes results in predictable outcomes

We next asked if network-rationalized interventions result in predictable outcomes upon perturbation, e.g., gene depletion (CRISPR, shRNA, KO mice) or overexpression, expression of functionally defective mutants, or chemical agonists/inhibitors. To this end, we carried out real-world *crowdsourcing experiments* on macrophage datasets in which interventions were conducted by different groups using diverse manipulations (**Fig 6a**). Depletion or pharmacologic inhibition of any gene in C#13 was predicted to suppress reactivity and enhance tolerance, whereas overexpression or pharmacologic stimulation of the same should have an opposite impact, i.e., enhance reactivity and suppress tolerance. Similarly, depletion/inhibition of any gene in C#14 was predicted to enhance reactivity and suppress tolerance (**Fig 6b**, *left*; **Table S3**). The depletion of genes in C#3 is predicted to not have a robust impact the network because of the Low=>Low relationship with C#14.

**Figure 6.**
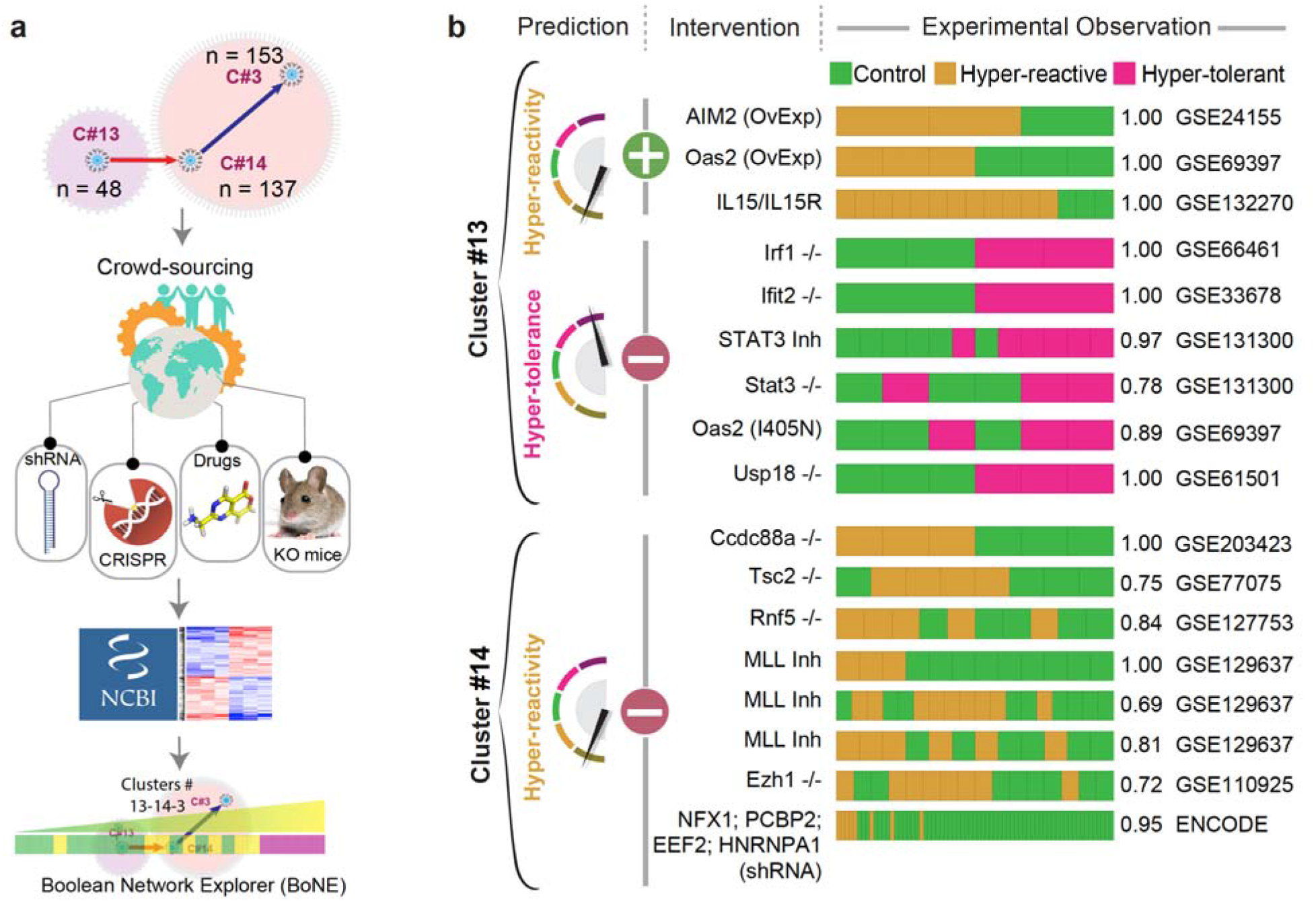
Crowd-sourced assessment of the predictive potential of the *SMaRT* genes. **a)** Overview of our workflow and approach for crowd-sourced validation. Publicly available transcriptomic datasets reporting the outcome of intervention studies (genetic or pharmacologic manipulations) on macrophages/monocytes targeting any of the 185 genes in C#13 and C#14 were analyzed using the *BoNE* platform for macrophage states. **b)** Predicted impact of positive (+, either overexpression [OvExp] or agonist stimulations) or negative (-; genetic -/-models, shRNA or chemical inhibitors) interventions and observed macrophage polarization states are shown. Performance is measured by computing *ROC AUC* for a logistic regression model. See **Table S3**.

We began with the ENCODE portal ^60^, a resource that was born out of the larger initiative called the ENCODE integrative analysis ^61^; it is an encyclopedia of large, unbiased shRNA library screen on the human *K562* chronic myeloid leukemia cell line. This dataset contained 4 of the 137 genes in C#14 and none from C#13 ^60^. In all 4 cases, the depletion of genes in C#14 resulted in the predicted outcome of enhanced reactivity and hypotolerance (**Fig 6b**, *right*). A systematic search of the NCBI GEO database also revealed 16 other independent datasets reporting the impact of interventions on genes in C#13 (9 datasets) and C#14 (7 datasets) (**Table S3**). Regardless of the heterogeneous nature of the interventions and lab-to-lab variations in the type of cells/tissues used, predictions matched the observed outcomes in each instance. At least in one instance (i.e., STAT3), we could confirm the alignment of phenotypes between gene deletion and pharmacologic inhibition, implying that both approaches must have converged on the same biology. Because such alignment and/or convergence is seen in many instances ^62^, findings suggest that the current model can accurately guide outcome-driven pharmacologic interventions.

Together, these crowd-sourced studies rigorously and independently validate the definitions of macrophage polarization states; the fundamental nature of these definitions appear to remain relevant despite the thunderous heterogeneity of models and methods used by so many.

## CONCLUSIONS

The lack of consensus on how to define macrophage activation has impeded progress in multiple ways; despite a panoply of existing descriptors, most remain contentious and/or confusing. AI-guided gene expression signatures presented here, *SMaRT*, offers a set of standardized definitions of macrophage polarization that encompasses four principles: (i) they are comprised of an unbiased collection of markers of macrophage activation that are represented in both the transcriptome and the proteome; (ii) they remain meaningful and relevant regardless of the source of macrophages (i.e., bone marrow, circulation, tissue-resident); (iii) they perform well across diverse activators, both in vitro and in vivo (i.e., recombinant ligands and cytokines, microbes, or multifactorial, as in the setting of complex disease states), and (iv) they provide a predictive framework that can be exploited for diagnostic purposes and for outcome-rationalized therapeutic interventions. These principles unify experimental standards for diverse experimental scenarios and interpretations across diverse tissues and diseases.

Finaly, these *SMaRT* genes provide a common framework for macrophage activation nomenclature, which should enable laboratories to detect and report a given immunophenotype of macrophage in a standardized way. Standardization is expected to spur the development of robust strategies to address the multitude of macrophage-related disorders. It also serves as a starting point for the development of new diagnostics and immunomodulatory therapies.

## Acknowledgements

We thank Gordon Gill and Christopher K. Glass for critiques and suggestions during manuscript preparation. This work was supported by the National Institutes for Health (NIH) grant R01-AI155696 (to PG, DS and SD). Other sources of support include: R01-GM138385 (to DS), R01-AI141630, CA100768 and CA160911 (to P.G), R01-DK107585 (to S.D) and UG3TR002968 (to D.S., S.D and P.G). D.S was also supported by two Padres Pedal the Cause awards (Padres Pedal the Cause/RADY #PTC2017 and San Diego NCI Cancer Centers Council (C3) #PTC2017). D.S, P.G and S.D were also supported by the Leona M. and Harry B. Helmsley Charitable Trust.

## Author Contributions

Conceptualization: D.S, P.G.

Methodology: D.S, S.S, D.V., S.T., D.D.

Investigation: D.S, S.S, P.G

Visualization: D.S, P.G, S.S, G.D.K, D.V.

Funding acquisition: D.S, S.D, P.G

Project administration: D.S, P.G

Supervision: D.S, P.G

Writing – original draft: D.S, P.G

Writing – review & editing: D.S, P.G, S.D, G.D.K, S.S

## Data and materials availability

All data are available in the main text or the supplementary materials. A website (http://hegemon.ucsd.edu/SMaRT/) of the macrophage network is built to support interactive query.

