## Supplemental Online Materials for "Machine Learning Identifies Signatures of Macrophage Reactivity and Tolerance that Predict Disease Outcomes"

**This PDF file includes:**

Materials and Methods

Figure S1 to S7

Table S1 to S3

**Materials and Methods**

**Key resources table**

| **REAGENT or RESOURCE** | **SOURCE** | **IDENTIFIER** |
| --- | --- | --- |
| **Deposited data** | | |
| Pooled human macrophage array | NCBI GEO (The National Center for Biotechnology Information- Gene expression omnibus) | [GSE134312](https://www.ncbi.nlm.nih.gov/geo/query/acc.cgi?acc=GSE134312) |
| *Ccdc88a* KO peritoneal macrophages |  | [GSE203423](https://www.ncbi.nlm.nih.gov/geo/query/acc.cgi?acc=GSE203423) |
| Proteomics dataset, reanalyzed from PMID: 34731634 | MassIVE repository | [MSV000084672](https://massive.ucsd.edu/ProteoSAFe/dataset.jsp?accession=MSV000084672) |
| **Experimental models**: Organisms/strains | | |
| *Ccdc88a fl/fl LysMCre/- mice* | PMID: 33055214 |  |
| **Software and algorithms** | | |
| Numpy | Python | <https://numpy.org> |
| Scipy | Python | <https://scipy.org> |
| Seaborn | Python | <https://seaborn.pydata.org> |
| Matplotlib | Python | <https://matplotlib.org> |
| Hierarchical Exploration of Gene Expression Microarrays Online (Hegemon) | HTML, JavaScript, Python, PHP | <https://github.com/sahoo00/Hegemon> |
| Boolean Network Explorer (BoNE) | Python | <https://github.com/sahoo00/BoNE> |
| Other | | |
| Interactive website | This paper | <http://hegemon.ucsd.edu/SMaRT/> |

**Data Collection and Annotation**

Publicly available microarray and RNASeq databases were downloaded from the National Center for Biotechnology Information (NCBI) Gene Expression Omnibus (GEO) website (Barrett et al. 2005; Barrett et al. 2013; Edgar et al. 2002). Gene expression summarization was performed by normalizing Affymetrix platforms by RMA (Robust Multichip Average) (Irizarry et al. 2003a; Irizarry et al. 2003b) and RNASeq platforms by computing TPM (Transcripts Per Millions) (Li and Dewey 2011; Pachter 2011) values whenever normalized data were not available in GEO. We used log2(TPM) if TPM > 1 and (TPM – 1) if TPM < 1 as the final gene expression value for analyses. We also used log2(TPM + 1) in some datasets. We also used publicly data normalized using RPKM (Mortazavi et al. 2008), FPKM (Trapnell et al. 2009; Trapnell et al. 2010), TPM (Li et al. 2010; Wagner et al. 2012), and CPM (Law et al. 2016; Robinson et al. 2010). In the context of Affymetrix microarray data we believe that RMA works better than MAS 5.0 (Pandey and Sahoo 2019).

*Macrophage datasets used for network analysis*

Previously published pooled macrophage dataset from GEO (GSE134312, n = 197) assayed on the Human U133 Plus 2.0 (GPL570), Human U133A 2.0 (GPL571) and Human U133A (GPL96) platforms were used to perform macrophage network analysis. This dataset was manually annotated with M0, M1 or M2 phenotypes. Accession numbers for the M0, M1 and M2 phenotypes are presented in table S4. Five validation datasets are used to test the macrophage gene signature: GSE35449 (7 M0, 7 M1, 7 M2), GSE46903 (64 M0, 29 M1, 40 M2), GSE61298 (6 M0, 6 M1, 6 M2), GSE55536 human peripheral blood mononuclear cell-derived macrophage (6 M0, 6 M1, 6 M2), GSE55536 iPSC derived macrophages (3 M0, 3 M1, 3 M2). See **Supplementary Information 1** for all datasets analyzed in this work.

**Computational Approaches**

*StepMiner Analysis*

StepMiner is a computational tool that identifies step-wise transitions in a time-series data (Sahoo et al. 2007). StepMiner performs an adaptive regression scheme to identify the best possible step up or down based on sum-of-square errors. The steps are placed between time points at the sharpest change between low expression and high expression levels, which gives insight into the timing of the gene expression-switching event. To fit a step function, the algorithm evaluates all possible step positions, and for each position, it computes the average of the values on both sides of the step for the constant segments. An adaptive regression scheme is used that chooses the step positions that minimize the square error with the fitted data. Finally, a regression test statistic is computed as follows:

$$F stat= \frac{{\sum_{i=1}^{n} {(\hat{X_{i}} - \bar{X})}^{2}}/{(m-1)}}{{\sum_{i=1}^{n} {(X_{i}- \hat{X_{i}})}^{2}}/{(n-m)}}$$

Where $X_{i}$ for$i=1$ to$n$ are the values, $\hat{X_{i}}$ for$i=1$ to$n$ are fitted values. m is the degrees of freedom used for the adaptive regression analysis. $\bar{X}$ is the average of all the values: $\bar{X}= \frac{1}{n}* \sum_{j=1}^{n} X_{j}.$For a step position at k, the fitted values $\hat{X_{l}}$ are computed by using $\frac{1}{k}* \sum_{j=1}^{n} X_{j}$ for$i=1$ to$k$ and $\frac{1}{(n-k)}* \sum_{j=k+1}^{n} X_{j}$ for$i=k+1$ to$n$.

*Boolean Analysis*

**Boolean logic** is a simple mathematic relationship of two values, i.e., high/low, 1/0, or positive/negative. The Boolean analysis of gene expression data requires the conversion of expression levels into two possible values. The ***StepMiner*** algorithm is reused to perform Boolean analysis of gene expression data (Sahoo et al. 2008). **The Boolean analysis** is a statistical approach which creates binary logical inferences that explain the relationships between phenomena. Boolean analysis is performed to determine the relationship between the expression levels of pairs of genes. The ***StepMiner*** algorithm is applied to gene expression levels to convert them into Boolean values (high and low). In this algorithm, first the expression values are sorted from low to high and a rising step function is fitted to the series to identify the threshold. Middle of the step is used as the StepMiner threshold. This threshold is used to convert gene expression values into Boolean values. A noise margin of 2-fold change is applied around the threshold to determine intermediate values, and these values are ignored during Boolean analysis. In a scatter plot, there are four possible quadrants based on Boolean values: (low, low), (low, high), (high, low), (high, high). A Boolean implication relationship is observed if any one of the four possible quadrants or two diagonally opposite quadrants are sparsely populated. Based on this rule, there are six kinds of Boolean implication relationships. Two of them are symmetric: equivalent (corresponding to the positively correlated genes), opposite (corresponding to the highly negatively correlated genes). Four of the Boolean relationships are asymmetric, and each corresponds to one sparse quadrant: (low => low), (high => low), (low => high), (high => high). BooleanNet statistics (**Fig. 2a**) is used to assess the sparsity of a quadrant and the significance of the Boolean implication relationships (Sahoo et al. 2008; Sahoo et al. 2010). Given a pair of genes A and B, four quadrants are identified by using the StepMiner thresholds on A and B by ignoring the Intermediate values defined by the noise margin of 2 fold change (+/- 0.5 around StepMiner threshold). Number of samples in each quadrant are defined as a_00_, a_01_, a_10_, and a_11_ (Figure 1A) which is different from X in the previous equation of F stat. Total number of samples where gene expression values for A and B are low is computed using the following equations.

${nA}_{low}= \left( a_{00}+ a_{01} \right), {nB}_{low}= \left( a_{00}+ a_{10} \right)$*,*

Total number of samples considered is computed using following equation.

$total= a_{00}+ a_{01}+ a_{10}+ a_{11}$

Expected number of samples in each quadrant is computed by assuming independence between A and B. For example, expected number of samples in the bottom left quadrant e_00_ = $\hat{n}$ is computed as probability of A low ((a_00_ + a_01_)/total) multiplied by probability of B low ((a_00_ + a_10_)/total) multiplied by total number of samples. Following equation is used to compute the expected number of samples.

$n= a_{ij}$*,* $\hat{n}= \left( {{nA}_{low}}/{total}* {{nB}_{low}}/{total} \right)*total$

To check whether a quadrant is sparse, a statistical test for (e_00_ > a_00_) or ($\hat{n}>n)$ is performed by computing S_00_ and p_00_ using following equations. A quadrant is considered sparse if S_00_ is high ($\hat{n}>n)$ and p_00_ is small.

$$S_{ij}= \frac{\hat{n}-n}{\sqrt{\hat{n}}}$$

$$p_{00}= \frac{1}{2} \left( \frac{a_{00}}{(a_{00}+ a_{01})}+ \frac{a_{00}}{(a_{00}+a_{10})} \right)$$

A suitable threshold is chosen for S_00_ > sThr and p_00_ < pThr to check sparse quadrant. A Boolean implication relationship is identified when a sparse quadrant is discovered using following equation.

***Boolean Implication*** = (*S_ij_* > sThr, *p_ij_* < pThr)

A relationship is called Boolean equivalent if top-left and bottom-right quadrants are sparse.

*Equivalent* $= \left( S_{01}> sThr, P_{01}< pThr, S_{10}> sThr, P_{10}< pThr \right)$

Boolean opposite relationships have sparse top-right (a_11_) and bottom-left (a_00_) quadrants.

*Opposite*$= \left( S_{00}> sThr, P_{00}< pThr, S_{11}> sThr, P_{11}< pThr \right)$

Boolean equivalent and opposite are symmetric relationship because the relationship from A to B is same as from B to A. Asymmetric relationship forms when there is only one quadrant sparse (A low => B low: top-left; A low => B high: bottom-left; A high=> B high: bottom-right; A high => B low: top-right). These relationships are asymmetric because the relationship from A to B is different from B to A. For example, A low => B low and B low => A low are two different relationships.

A low => B high is discovered if the bottom-left (a_00_) quadrant is sparse and this relationship satisfies following conditions.

*A low => B high* = ($S_{00}> sThr, P_{00}< pThr$)

Similarly, A low => B low is identified if the top-left (a_01_) quadrant is sparse.

*A low => B low* = ($S_{01}> sThr, P_{01}< pThr$)

A high => B high Boolean implication is established if the bottom-right (a_10_) quadrant is sparse as described below.

*A high => B high* = ($S_{10}> sThr, P_{10}< pThr$)

Boolean implication A high => B low is found if the top-right (a_11_) quadrant is sparse using following equation.

*A high => B low* = ($S_{11}> sThr, P_{11}< pThr$)

For each quadrant a statistic S_ij_ and an error rate p_ij_ is computed. S_ij_ > sThr and p_ij_ < pThr are the thresholds used on the BooleanNet statistics to identify Boolean implication relationships.

Boolean analyses in the test dataset GSE134312 uses a threshold of sThr = 3 and pThr = 0.1. These thresholds are exactly same as the previously used thresholds sThr = 3 and pThr = 0.1 for BooleanNet (Dabydeen et al. 2019; Pandey and Sahoo 2019; Sahoo et al. 2008).

*Boolean Network Explorer (BoNE)*

Boolean network explorer (BoNE) provides an integrated platform for the construction, visualization and querying of a network of progressive changes underlying a disease or a biological process in three steps (**Fig S1A**): First, the expression levels of all genes in these datasets were converted to binary values (high or low) using the StepMiner algorithm. Second, gene expression relationships between pairs of genes were classified into one-of-six possible Boolean Implication Relationships (BIRs), two symmetric and four asymmetric, and expressed as Boolean implication statements. This offers a distinct advantage from conventional computational methods (Bayesian, Differential, etc.) that rely exclusively on symmetric linear relationships in networks. The other advantage of using BIRs is that they are robust to the noise of sample heterogeneity (i.e., healthy, diseased, genotypic, phenotypic, ethnic, interventions, disease severity) and every sample follows the same mathematical equation, and hence is likely to be reproducible in independent validation datasets. Third, genes with similar expression architectures, determined by sharing at least half of the equivalences among gene pairs, were grouped into clusters and organized into a network by determining the overwhelming Boolean relationships observed between any two clusters. In the resultant Boolean implication network, clusters of genes are the nodes, and the BIR between the clusters are the directed edges; BoNE enables their discovery in an unsupervised way while remaining agnostic to the sample type.

*Statistical Analyses*

Gene signature is used to classify sample categories and the performance of the multi-class classification is measured by ROC-AUC (Receiver Operating Characteristics Area Under The Curve) values. A color-coded bar plot is combined with a density or violin+swarm plot to visualize the gene signature-based classification. All statistical tests were performed using R version 3.2.3 (2015-12-10). Standard t-tests were performed using python scipy.stats.ttest_ind package (version 0.19.0) with Welch’s Two Sample t-test (unpaired, unequal variance (equal_var=False), and unequal sample size) parameters. Multiple hypothesis corrections were performed by adjusting *p* values with statsmodels.stats.multitest.multipletests (fdr_bh: Benjamini/Hochberg principles). The results were independently validated with R statistical software (R version 3.6.1; 2019-07-05). Pathway analysis of gene lists were carried out via the Reactome database and algorithm (Fabregat et al. 2018). Reactome identifies signaling and metabolic molecules and organizes their relations into biological pathways and processes. Kaplan-Meier analysis is performed using lifelines python package version 0.14.6.

*Boolean implication network construction*

A Boolean implication network (BIN) is created by identifying all significant pairwise Boolean implication relationships (BIRs) for GSE134312 datasets (**Fig S1A**). The Boolean implication network contains the six possible Boolean relationships between genes in the form of a directed graph with nodes as genes and edges as the Boolean relationship between the genes. The nodes in the BIN are genes and the edges correspond to BIRs. Equivalent and Opposite relationships are denoted by undirected edges and the other four types (low => low; high => low; low => high; high => high) of BIRs are denoted by having a directed edge between them. The network of equivalences seems to follow a scale-free trend; however, other asymmetric relations in the network do not follow scale-free properties. BIR is strong and robust when the sample sizes are usually more than 200. However, it is also possible to build BIN for smaller dataset such as the selected macrophage GSE134312 dataset (n = 197). The macrophage dataset GSE134312 was prepared for Boolean analysis by filtering genes that had a reasonable dynamic range of expression values. When the dynamic range of expression values was small, it was difficult to distinguish if the values were all low or all high or there were some high and some low values. Thus, it was determined to be best to ignore them during Boolean analysis. The filtering step was performed by analyzing the fraction of high and low values identified by the StepMiner algorithm (Sahoo et al. 2007). Any probe set or genes which contained less than 5% of high or low values were dropped from the analysis.

*Clustered Boolean Implication network*

Clustering was performed in the Boolean implication network to dramatically reduce the complexity of the network (**Fig S1C**). A clustered Boolean implication network (CBIN) was created by clustering nodes in the original BIN by following the equivalent BIRs. One approach is to build connected components in a undirected graph of Boolean equivalences. However, because of noise the connected components become internally inconsistent e.g. two genes opposite to each other becomes part of the same connected component. In order to avoid such situation, we need to break the component by removing the weak links. To identify the weakest links, we first computed a minimum spanning tree for the graph and computed Jaccard similarity coefficient for every edge in this tree. Ideally if two members are part of the same cluster they should share as many connections as possible. If they share less than half of their total individual connections (Jaccard similarity coefficient less than 0.5) the edges are dropped from further analysis. Thus, many weak equivalences were dropped using the above algorithm leaving the clusters internally consistent. We removed all edges that have Jaccard similarity coefficient less than 0.5 and built the connected components with the rest. The connected components were used to cluster the BIN which is converted to the nodes of the CBIN. The distribution of cluster sizes was plotted in a log-log scale to observe the characteristic of the Boolean network (**Fig S1C**). The clusters sizes were distributed along a straight line in a log-log plot suggesting scale-free properties (**Fig S1D**). A new graph was built that connected the individual clusters to each other using Boolean relationships. Link between two clusters (A, B) was established by using the top representative node from A that was connected to most of the member of A and sampling 6 nodes from cluster B and identifying the overwhelming majority of BIRs between the nodes from each cluster.

A CBIN was created using the selected GSE134312 datasets. Each cluster was associated with healthy or disease samples based on where these gene clusters were highly expressed. The edges between the clusters represented the Boolean relationships that are color-coded as follows: orange for low => high, dark blue for low => low, green for high => high, red for high => low, light blue for equivalent and black for opposite.

*Boolean paths*

The asymmetric BIRs provide a unique dimension to the network that is fundamentally different from any other gene expression networks in the literature. Traversing a set of nodes in a directed graph of the Boolean network constitutes a Boolean path that can be interpreted as follows. A simple Boolean path involves two nodes and the directed edge between them. This simple Boolean path can be interpreted as shown in the supplementary figure (**Fig S1E**). For the nodes X and Y with X low => Y low only quadrant #1 is sparse; the other quadrants #0, #2, and #3 are filled with samples (**Fig S1E**). Assuming monotonicity in X and Y, the quadrants can be ordered in two possible ways: 0-2-3 and 3-2-0. The path corresponds to 0-2-3 begins with X low and Y low. This is interpreted as X turns on first and then Y turns on along a hypothetical biological path defined by the sample order. Similarly, Y turns off first and then X turns off in the path 3-2-0. A complex path in the Boolean network involves more than one Boolean implication relationship (**Fig S1F**). Three Boolean implication relationships can be used to group samples into five bins and the bins can be ordered in two possible ways (**Fig S1F**, forward, reverse). Another example of a path is illustrated in supplementary figure (**Fig S1G**).

*Discovery of Paths in Clustered Boolean Implication network*

Discovery of paths start with a node that represents the biggest cluster in the CBIN. Since a path of high=>high, high=>low, and low=>low can be used to order samples as shown in **Fig S1G**, we try to identify paths of this type that intersects the big clusters in the network. We developed a simple, intuitive algorithm that traverses the nodes of the CBIN starting with the biggest cluster and greedily chooses next big cluster connected to the nodes visited in sequence. The emphasis on cluster sizes comes from the fundamental assumption that size determines importance and relevance. Therefore, we start from a big cluster (A1) and identify other clusters that form a chain of low => low. Further, we identify other clusters that are either opposite to A1 or they have high=>low relationship with A1, and the biggest cluster (A2) among these clusters were chosen. In addition, a chain of low=> low relationship from A2 is identified. In each subsequent step, again the biggest cluster among the different choices was greedily chosen. Finally equivalence relationship from each cluster is used to gather more genes in each cluster and the whole path is clustered based on equivalence relationships. Depth-first traversal (DFS) was used to follow the path of low => low where bigger clusters are visited first. The search was performed until a cluster was reached for which there is no low => low relationships. For example, starting with cluster S, the search will return S low => A1 low, A1 low => A2 low, and A2 low => A3 low if A3 doesn’t have any low => low relationships. Similarly, a new starting point is considered S2 such that S2 is the biggest cluster X that has either S high => X low or S Opposite X. From cluster S2 another DFS was performed to retrieve the longest possible path of low => low. The search may return S2 low => B1 low, B1 low => B2 low if B2 doesn’t have any low => low relationships. In summary, the most prominent Boolean path was discovered by starting with the largest cluster and then exploring edges that connected to the next largest cluster in a greedy manner. This process was repeated to explore paths that connect the big clusters in the network.

*Scoring Boolean path for sample order*

A score was computed for a specified Boolean path that can be used to order the sample which was consistent with the logical order. To compute the final score, first the genes present in each cluster were normalized and averaged. Gene expression values were normalized according to a modified Z-score approach centered around StepMiner threshold (formula = (expr - SThr)/3*stddev; **Fig S2B**). Weighted linear combination of the averages from the clusters of a Boolean path was used to create a score for each sample. The weights along the path either monotonically increased or decreased to make the sample order consistent with the logical order based on BIR. The samples were ordered based on the final weighted (-1 for C#13, 1 for C#14 and 2 for C#3) and linearly combined score (**Fig S2C**). The direction of the path was derived from the connection from a reactive cluster to a tolerant cluster. The sample order is visualized by a color-coded bar plot and a violin+swarm plot (**Fig S2C**).

*Summary of genes in the clusters*

Reactome pathway analysis of each cluster along the top continuum paths was performed to identify the enriched pathways (Fabregat et al. 2018). The pathway description was used to summarize at a high-level what kind of biological processes are enriched in a particular cluster.

*Signatures of macrophage reactivity and tolerance (S-Ma-R-T) computation*

*BoNE* uses Boolean implication network on macrophage dataset to build a signature of macrophage polarization. Selected clusters by size connected by high => high (green arrow), high => low (red arrows) and low => low (blue arrows) Boolean implication relationships. Reactome analysis of each clusters shows the biological processes the genes are involved in (**Fig S2A**). A path is selected in the network that is used to test M1/M2 states classification. This process is demonstrated by using a path #13-14-3 on GSE134312 (**Fig S2B-C**).

*Single cell data analysis*

Single cell datasets were processed using scanpy (v1.5.1) framework. Composite scores for C#13 (weight = -1) and C#14-3 (weights = 1, 2) were computed like bulk RNASeq datasets. Scatterplot between C13 and C14-3 score were plotted using pandas plotting functions. A box is drawn manually (in the lower middle corner) that enrich reactive macrophages based on the data from controlled in-vitro reprogramming experiments.

*Normalization of gene expression based on circadian rhythm*

Since the state of macrophage swings from reactive to tolerant from day to night, it is important to control for this variation during analysis of macrophage polarization. To start the normalization process, clock genes (such as DBP, ARNTL, etc.) or gene signatures that capture circadian rhythm is used to adjust the BoNE score (**Fig S4**). First, both the BoNE score (**Fig S4B**) and the clock gene expression are scaled for each sample type based on their dynamic range of expression values (min – max). For example, the dataset GSE98895 contains two sample types: C (Control), and MetS (Metabolic Syndrome). Let’s take one sample from the MetS group (x, y) where x is the clock gene expression value and y is the original BoNE score (**Fig S4C**). Bounding box for the MetS group demonstrates the range of values for both the BoNE score (S1) and the clock gene expression (S2). Average of BoNE scores and the clock gene expression is shown using an orange diamond. The distance of (x, y) from the orange diamond (S3, S4) is used to scale both values: (x – S3 * (S2 + 1) / (S1 + 1) , y + S4 * (S1 + 1) /(S2 + 1)). This process is repeated using control (C) samples using the green diamond. Linear regression is used to compute the trend between the transformed BoNE score and clock gene expression (y = mx + c; **Fig S4D**). The trend is subtracted from the transformed BoNE score to compute the final normalized BoNE score (y – mx - c). Samples are now rank ordered based on the final normalized BoNE score to visualize the effect of normalization process.

*Proteomics analysis*

A multiplexed TMT (tandem mass tags) quantitative proteomics dataset has been obtained from He, L. et al (He et al. 2021) (see *Key Resource Table*). To generate this dataset, authors had differentiated human THP-1 cells with phorbol myristate acetate (PMA) for 24 h into macrophages (M0 state). The M0 cells were subsequently treated with IL4 for M2 polarization and with LPS and IFNγ for M1 polarization over a 24-h time-period. Samples were processed for quantitative mass spectrometry at 1 h, 4 h, 8 h and 24 h. Ratio of raw intensity values has been compared between M1 and M2 states to obtain the list of induced proteins at various time points (see **Supplemental Information 3**). To obtain the list of proteins induced in M1 state, the cut-off used for induction of proteins when comparing the raw intensity ratio for LPS/IFNγ over IL4 stimulation for all time points was >=2. To obtain the list of proteins induced in M2 state, the cut-off used for induction of proteins when comparing the raw intensity ratio for IL4 over LPS/IFNγ stimulation for all time points was >=1.5.

To assess the differential enrichment of proteins across different signatures for both M1 and M2 polarization states at various time points, we used the following equation to calculate the z-test of proportions,

z = $\frac{(p1-p2)}{\sqrt{p(1-p)\left( \frac{1}{n1}+\frac{1}{n2} \right)}}$

Here, p1 is sample proportion (x1/n1) of proteins translated from the “reactive” signature that were induced >=2 fold upon LPS stimulation. And p2 is the sample proportion (x2/n2) of proteins translated from the “tolerance” signature that were induced >=1.5 fold upon IL4 stimulation. Here, p = (x1+x2)/(n1+n2).

**Supplementary Figure Legends**

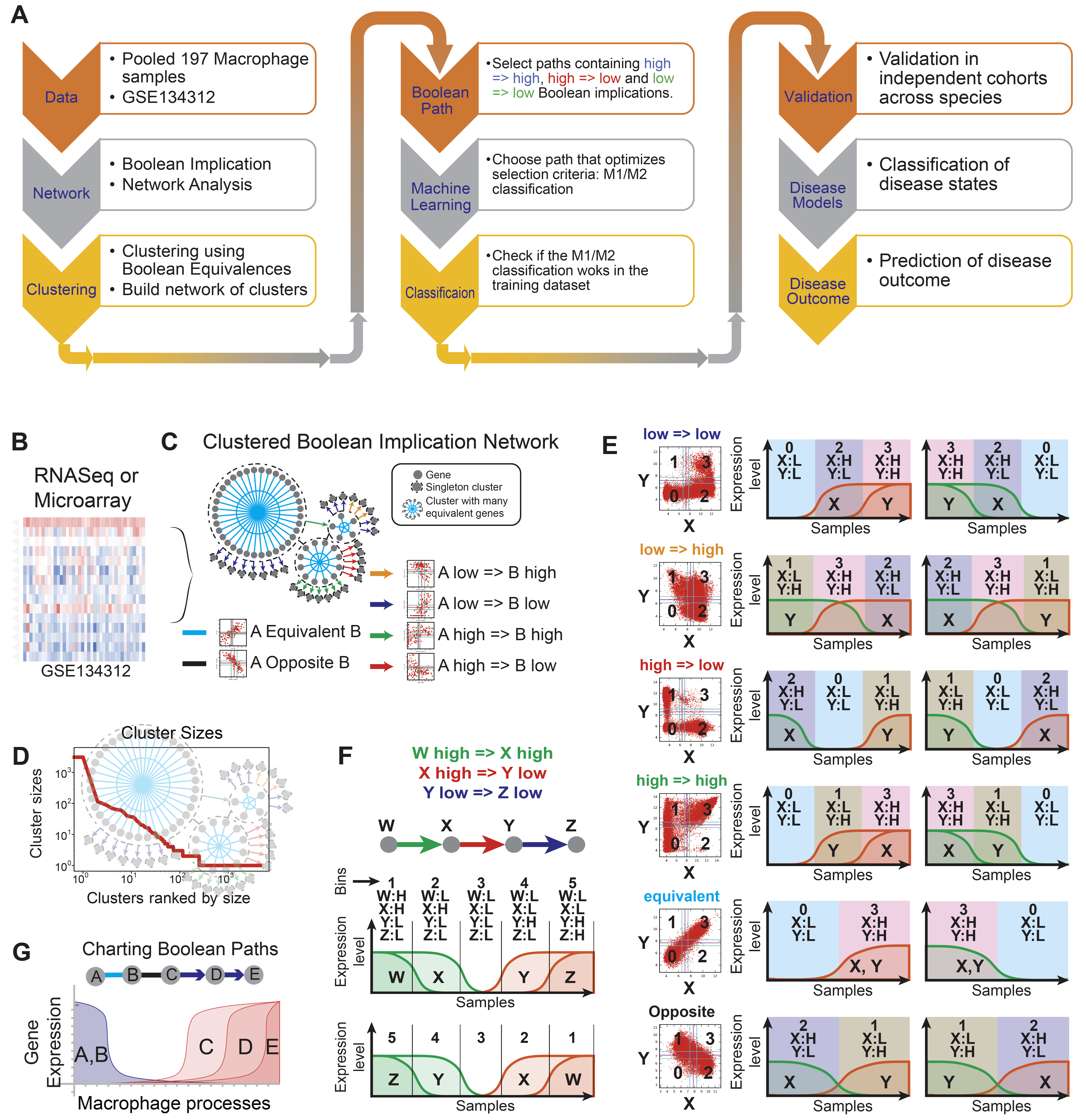

**Figure S1: Boolean Network Explorer (BoNE): A tool for clustering and visualization of the Boolean implication network**. **(A)** Overview of the computational steps used in *BoNE*. **(B)** *BoNE* was applied to analyze macrophage datasets to develop a model of polarization. GSE134312 is used to build the Boolean implication network. **(C)** BooleanNet algorithm is applied to identify Boolean implication relationships. The *BoNE* uses Boolean equivalent relationships to cluster genes and identify relationships between clusters. **(D)** A graphical display of cluster size analysis shows a linear trend in log-log scatterplots between clusters sorted by size and the number of clusters of any particular size. **(E)** Sample ordering based on single Boolean Implication relationships. **(F)** Sample ordering based on a sequence of high => high, high => low, low => low Boolean relationships. **(G)** Similar to panel F, four Boolean implication relationships ‘A equivalent to B’, ‘B opposite C’, ‘C low => D low’ and ‘D low => E’ low constitute a Boolean path that can be used to develop a computational model of macrophage polarization. A suitable path is selected by using machine learning that optimizes the strength of M1/M2 classification.

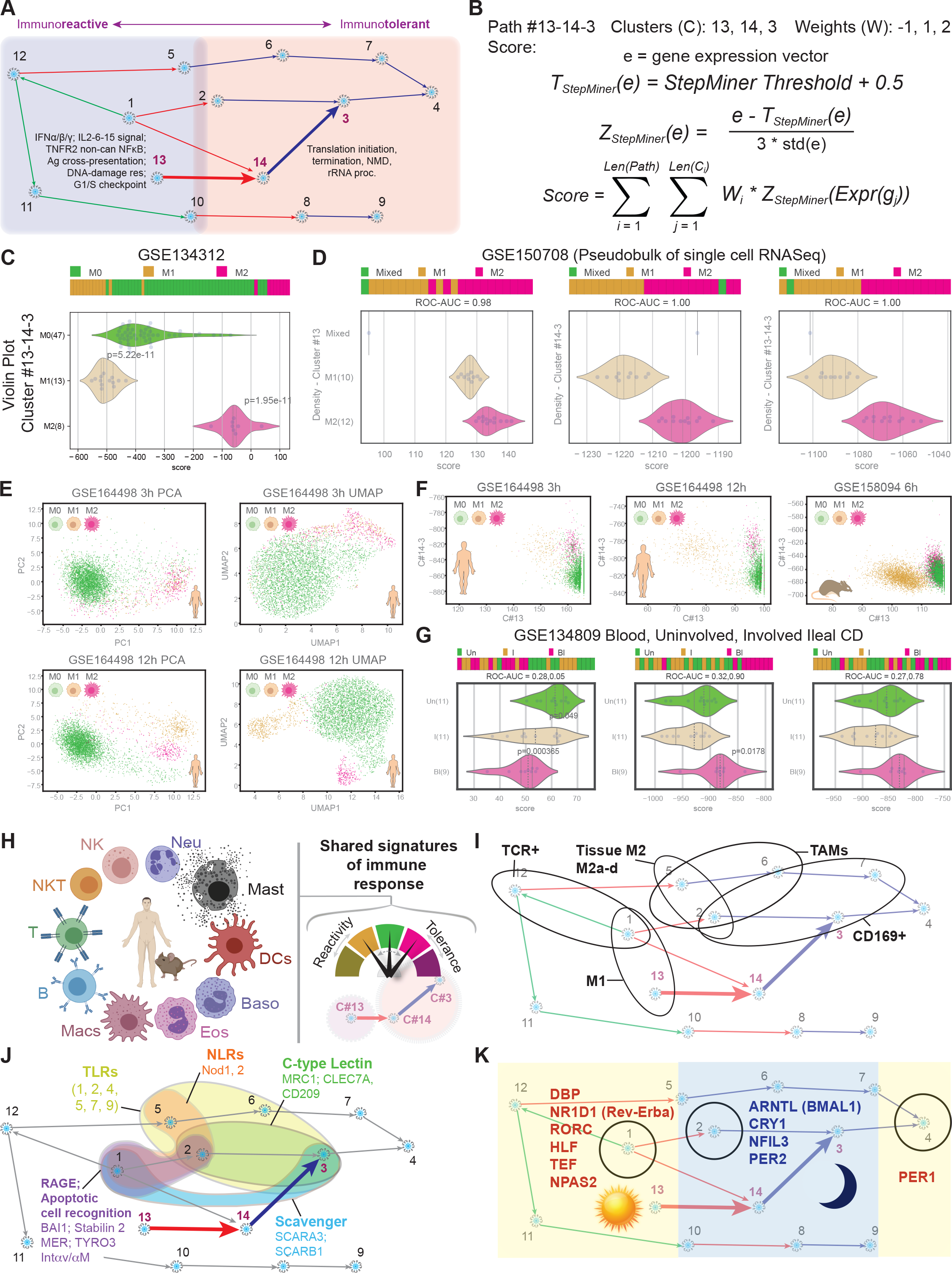

**Figure S2: Computational model of macrophage polarization: (A)** *BoNE* uses a Boolean implication network on macrophage dataset to build a computational model of macrophage polarization. Selected clusters by size were connected by high => high (green arrow), high => low (red arrows) and low => low (blue arrows) Boolean implication relationships. Reactome analysis of each cluster showed the biological processes in which the genes are involved.

**(B-C)** A path (#13-14-3) was selected in the network that is used to test M1/M2 states classification based on a score computed by using linear combination of normalized gene expression values (composite score). This process is demonstrated by using a path #13-14-3 on GSE134312. P-values are derived from Welch's two-sided unpaired unequal variance Two Sample t-test between control and experimental samples.

(**D**) Test of M1 and M2 classification using cluster #13, path #14-3, and path #13-14-3 using artificial mixture of M1/M2 macrophages in single cell RNASeq dataset GSE150708. Mixed sample was created by combining 4000 lung cells, 4000 M1 and 4000 M2 macrophages such that final pseudo-bulk sample contains 33% M1 and 33% M2 macrophages. ROC-AUC values of M1 vs M2 classification are shown below each bar plot. The rest of the samples contain 10% of different combination of M1 and M2 cells and 90% lung cells.

(**E**) PCA and UMAP analysis of human single cell dataset GSE164498 for 3h and 12h macrophage polarization protocols.

(**F**) Scatterplots of C#13 and path #14-3 composite scores in single cell RNASeq data GSE164498 (3h and 12h, human) and GSE158094 (6h, mouse).

(**G**) Bar and violin plots of pseudo-bulk analysis of computationally isolated macrophages (TYROBP > 2 and FCER1G > 2) from uninvolved and involved ileal biopsies and blood samples from Crohn’s disease patients.

(**H**) The schematic summarizes BoNE models (C#13, #14-3, #13-14-3) predicts activation states in diverse immune cell types.

(**I**) Known macrophage subtypes, as defined by marker genes, are projected on the Boolean map of macrophage processes. (**J**) The distribution of pattern recognition receptors (PRRs) [see **Table S2**] within various gene clusters of the Boolean map of macrophage processes is displayed.

(**K**) The positions of key circadian genes that are present in the network are shown on the Boolean map of macrophage processes. See also **Fig S4**.

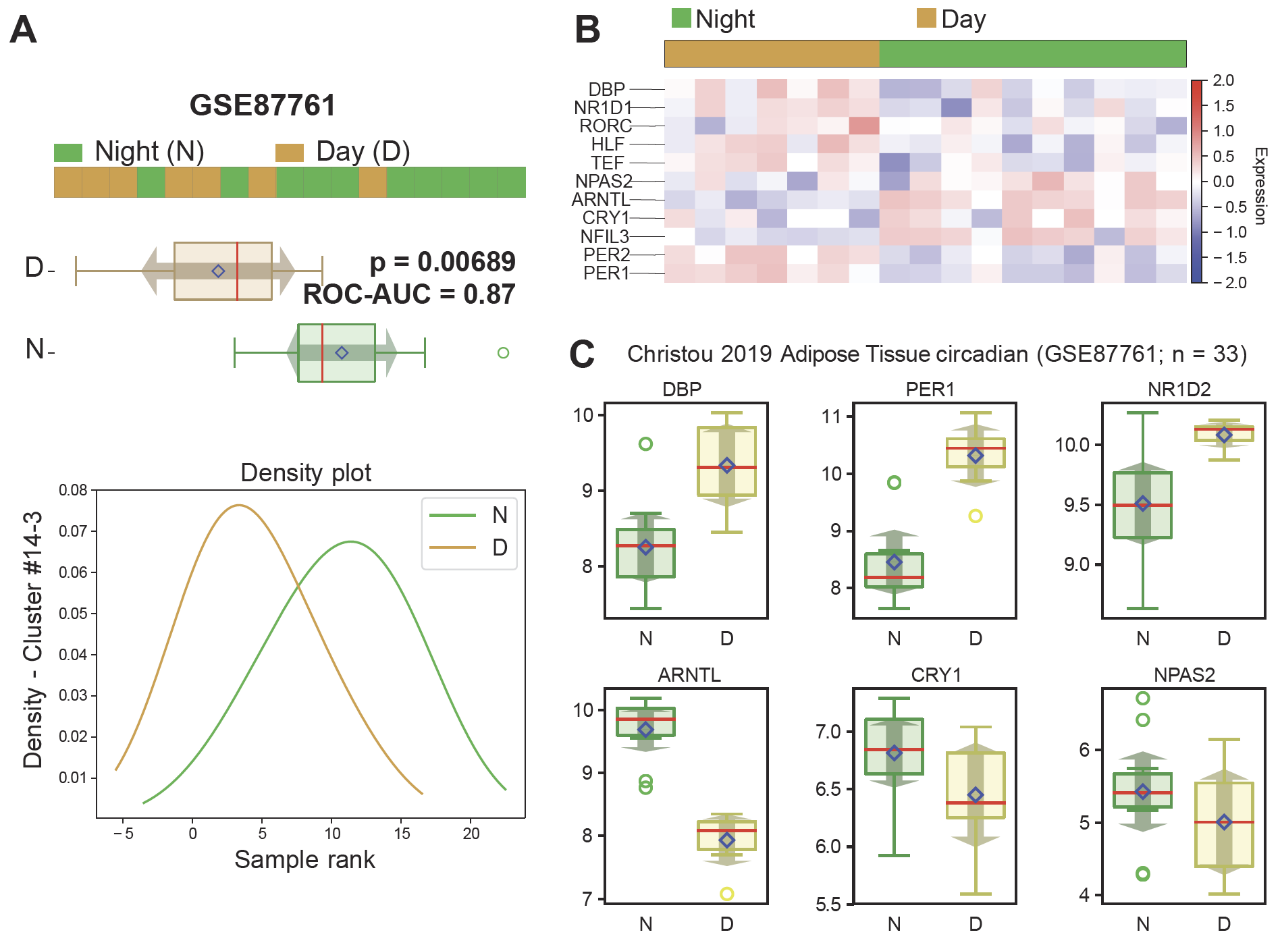

**Figure S3: Macrophages are reactive during the day and tolerant during the night.** **(A)**. Classification of macrophage polarization stages based on Boolean path #14-3 in GSE87761 (n=17). This dataset contains tissue samples from subcutaneous adipose tissue that were taken at regular intervals under the highly controlled conditions of a strict sleep/wake and meal schedule. Circadian phases from -4 to 6 relative to dim light melatonin onset (DLMO) is considered night (N) and from 6 to 12 is considered day (D). A barplot (top), a boxplot (middle) and a density plot (bottom) are shown to demonstrate the ordering of samples using *BoNE* score based on path #14-3. The density plot is based on the rank of the score whereas barplot and boxplot are based on the raw *BoNE* score (path #14-3). The results show that night (N) is more tolerant compared to daytime (D). P-values are derived from Welch's two-sided unpaired unequal variance Two Sample t-test between control and experimental samples. ROC-AUC values of N vs D classification are shown below the bar plot. **(B)** Expression of important circadian genes found in the macrophage network in D and N samples is displayed as heatmap. **(C)** Boxplots displaying the levels of expression of circadian genes found in the macrophage network.

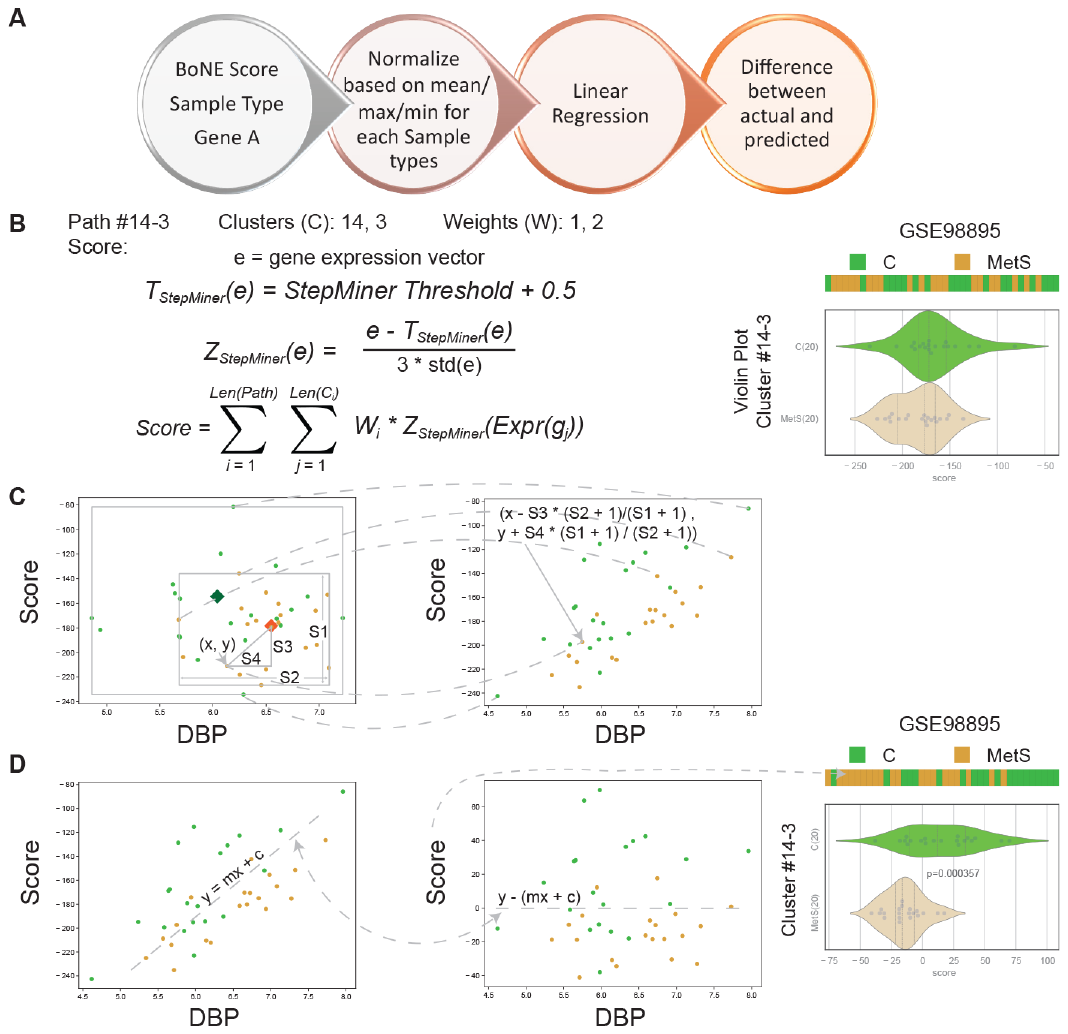

**Figure S4: Normalization of gene expression based on circadian rhythm:** **(A)** Overview of the normalization process. Normalization adjusts the *BoNE* score based on a clock gene (DBP, ARNTL, etc.). **(B)** Computation of original BoNE score based on a Boolean path. Example path #14-3 is used to demonstrate the score computation. StepMiner threshold + 0.5 is used to select the highest expression values. The expression values are scaled by using this threshold and the standard deviation. The *BoNE* score is computed by using a weighted linear combination of the scaled expression values. **(C)** Average values for each sample type are shown as colored diamonds. Maximum and minimum values are represented by bounding boxes. The *BoNE* score was adjusted based on maximum, minimum and average expressions of a clock gene in each sample type. The clock gene expression values were adjusted based on the *BoNE* score. **(D)** Linear regression was performed between the adjusted *BoNE* score and adjusted clock gene expression. Final normalized *BoNE* score was computed by subtracting the predicted trend. The final normalized score was used to rank the samples and visualized by using a bar+violin+swarm plot along with heatmap of sample types.

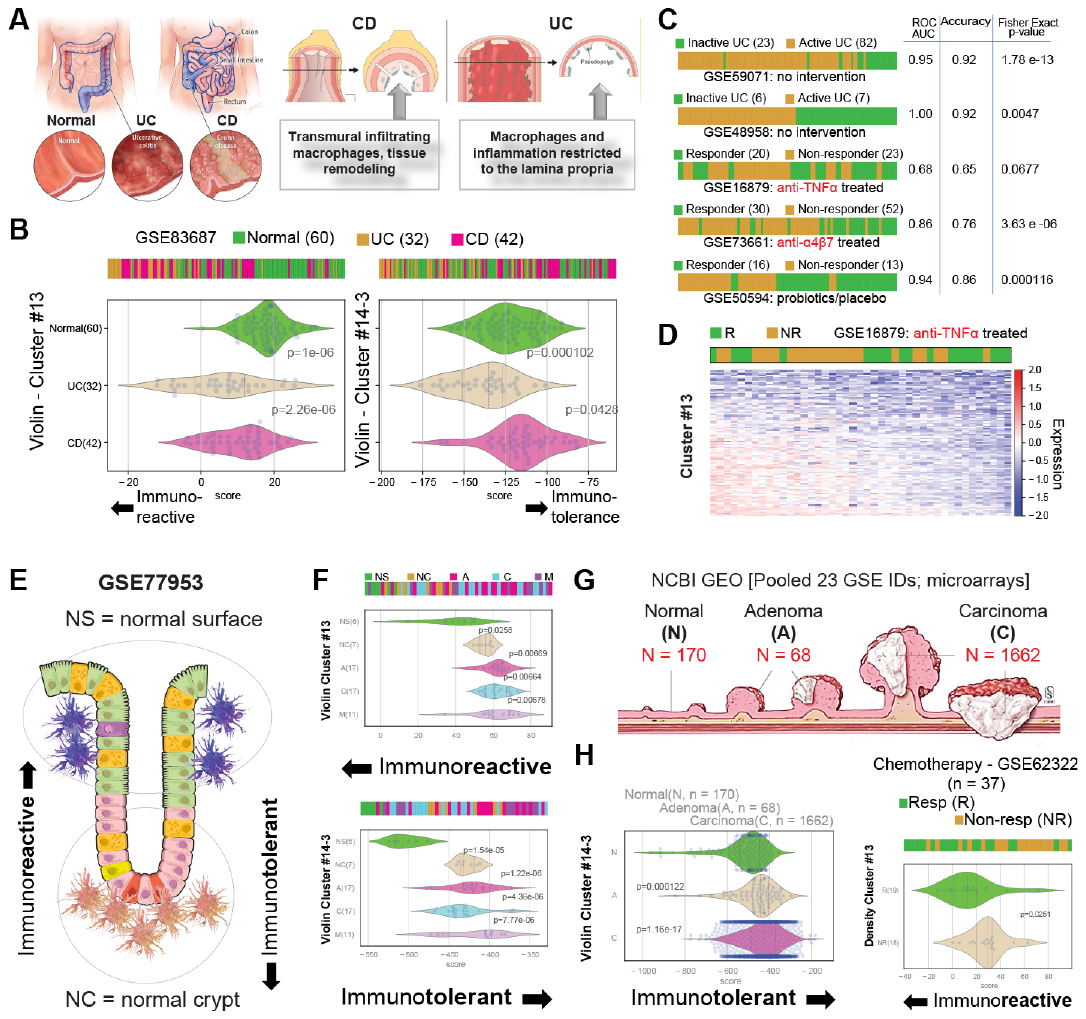

**Figure S5: Formal definitions of “reactivity” and “tolerance” identify the physiologic diversity of macrophages in the normal gut and the pathogenic responses in IBD and colorectal cancers (CRC).** (**A)** Normal, ulcerative colitis (UC) and crohn’s disease (CD) samples were analyzed based on cluster #13 and path #14-3 gene signatures, which revealed distinct disease states. **(B)** Analysis of an IBD RNA Seq dataset (GSE83687, n = 134, 60 N, 32 UC, 42 CD) revealed hyper-reactive macrophage states in both UC and CD, but hyper-tolerant macrophage state exclusively in CD. P-values are derived from Welch's two-sided unpaired unequal variance Two Sample t-test between control and disease samples. **(C)** Levels of expression of genes in cluster #13 can distinguish responders *vs*. non-responders to treatment with: anti-α4β7 (GSE73661, n=82, 30 R, 52 NR); anti-TNFα (GSE16879, n=24, 8 R, 16 NR); standard therapies comprised of steroids, mesalamine, etc. (GSE59071, n=97, 23 R, 74 NR; and GSE48958, n=13, 6 R, 7 NR); placebo/probiotic (GSE50594, n=29, 16 R, 13 NR). **(E)** Schematic summarizing the finding that the macrophages near the surface of normal crypt (NS) are more reactive than those near the base of normal crypt (NC). **(F)** Bar and violin plots show that Cluster 13 (top) and path 14-3 (bottom) gene signatures are differentially expressed in NS vs NC in laser-dissected tissues from the top and bottom of normal colon crypts and colon adenomas and CRCs (GSE77953, n=58, 6 NS, normal surface; 7 NC, normal crypt; 17 A, adenomas; 17 C, primary CRC; 11 M, metastatic CRC). (**G-H**) Cluster 13 and path 14-3 gene signatures analyzed on a pooled dataset (NCBI GEO dataset, 170 N, normal; 68 A, adenoma; 1662 C, Carcinoma) reveal a distinct progressive onset of tolerance in adenomas and CRCs compared to the normal colon (*left*) and that responders (R) to chemotherapy show higher reactivity than non-responders (NR) (GSE62322, n=37, 19 R, 18 NR) (*right*).

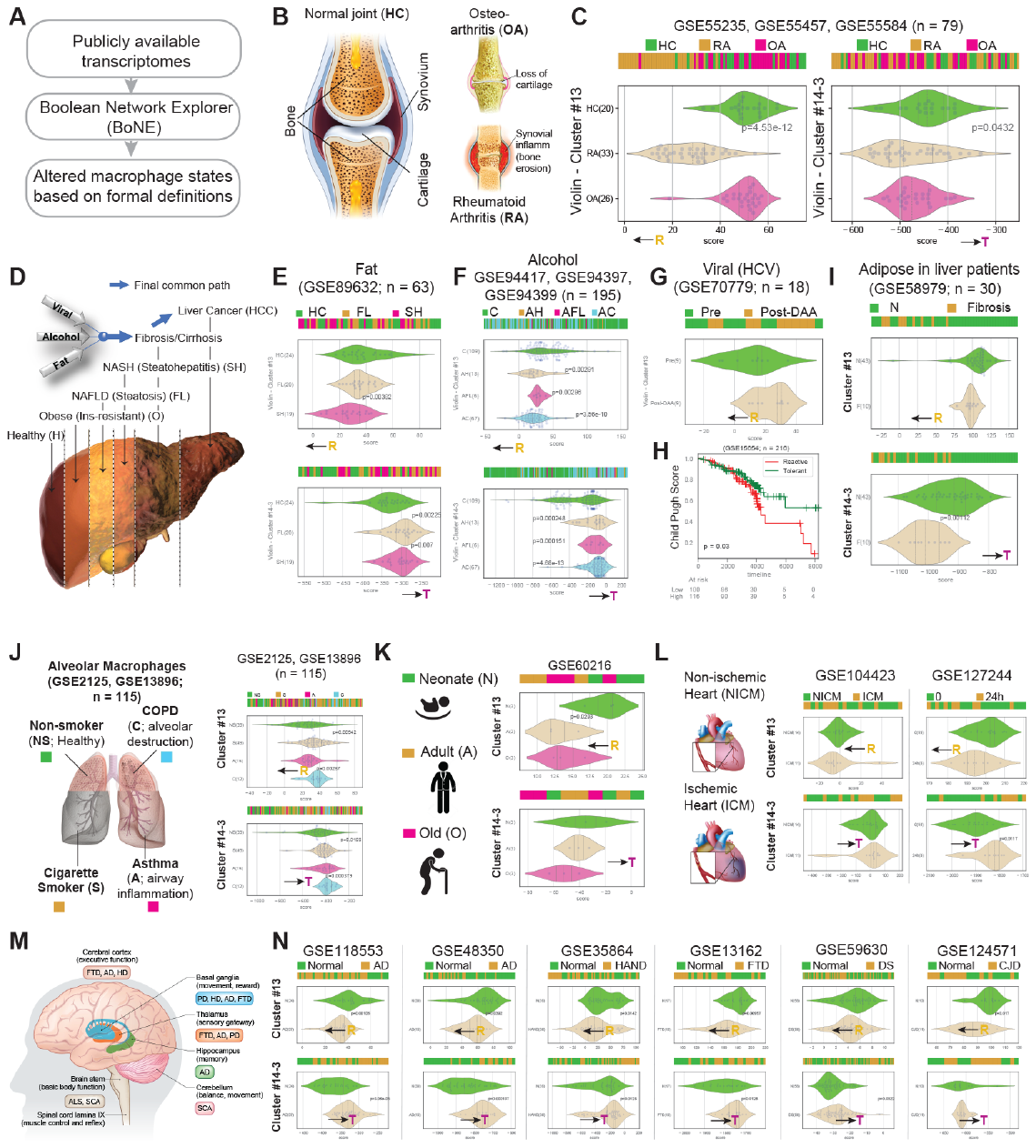

**Figure S6: Formal definitions of macrophage “reactivity” (R) and “tolerance” (T) identify pathologic states in diverse diseases.** (**A**) General approach used towards the analyses in **Figure 4, S6 and S7**. R denotes reactivity and ‘T’ denotes tolerance; the arrows show the direction of progressive reactivity and/or tolerance based on the levels of expression of gene signatures. P-values are derived from Welch's two-sided unpaired unequal variance Two Sample t-test between control and disease samples. (**B**) Summary of the two major types of arthritis-- rheumatoid arthritis (RA) involves swelling of synovium that may follow joint erosion, whereas osteoarthritis (OA) involves loss of cartilage between the joints. (**C**) Bar and violin plots display the analyses on three publicly available arthritis cohorts (GSE55235, GSE55457 and GSE55584; n = 79; 20 HC, 33 RA, 26 OA), which revealed that macrophages are hyper-reactive in RA whereas they are hypo-reactive in OA. However, with respect to tolerance, they are both similar to healthy controls (HC). (**D**) Multiple insults that trigger inflammation in the liver can cause progression to fibrosis (scarring), cirrhosis or liver cancer. (**E-G**) Bar and violin plots display the analyses on publicly available liver datasets from non-alcoholic fatty liver disease (**E**; GSE89632, n=63; 20 FL, fatty liver; 19 SH, steatohepatitis; 24 HC, healthy controls), pooled cohorts of patients with alcoholic liver disease (**F**; GSE94417, GSE94397, GSE94399, n = 195, 109 C, control; 13 AH, alcoholic hepatitis; 6 AFL, alcoholic fatty liver; 67 AC, alcoholic cirrhosis) and viral liver disease (**G**; GSE70779, n=18; 9 Pre- and 9 Post-treatment with direct-acting anti-viral [post-DAA]). (**H**) Among patients with Child-Pugh A cirrhosis, analysis of a prospective study (GSE15654, n = 216) showed that a higher reactive state (i.e., high expression of genes in cluster #13 *and* low expression of genes in #14-3) is unfavorable and is associated with a greater progression in Child-Pugh scores during follow-up. (**I**) Reactive state in subcutaneous and visceral fat is associated with liver fibrosis (F) as compared to NASH+NAFLD (N) (GSE58979, n = 53, 10 F, 43 N). (**J**) Schematic (*left*) summarizing the three major causes of the inflammation of the lung: smoking, asthma and chronic obstructive pulmonary disease (COPD); the latter is characterized by irreversible and progressive lung damage. Analysis (*right*) of two publicly available lung datasets (GSE2125, GSE13896, n = 115, 39 NS, non-smoker; 49 S, smoker; 15 A, asthma; 12, C, COPD) revealed that smoking and COPD are both hypo-reactive states (top) and that COPD is characterized by also hyper-tolerance (bottom). (**K**) Schematic (*left*) summarizing the three major age groups included to study the impact of aging on macrophage processes. Analysis of a publicly available dataset (*right*) from peripheral monocytes from three independent donors (GSE60216, n=9, 3 N, neonate; 3 A, adult; and 3 O, old adult) unstimulated across 3 time-points. (**L**) Schematic (*left*) shows the two different types of cardiomyopathies leading to heart failure, ischemic and non-ischemic. Analysis (*right*) of publicly available dataset from PBMCs from a cohort of heart failure patients (GSE104423; n = 25; 11 ICM, ischemic cardiomyopathy; 14 NICM, non-ischemic cardiomyopathy prior to undergoing mechanical circulatory support) and hearts of mice after experimental myocardial ischemia (GSE127244; n=16 at 0 and n=8 at 24 h) showed that ICM is associated with hyperreactivity and hypertolerance. (**M**) Schematic showing the various neurodegenerative diseases and the regions of the brain that they involve. (N) Analysis of multiple publicly available datasets representing a variety of neurodegenerative diseases. *From left to right*: AD, Alzheimer’s; HAND, HIV-associated neurodegenerative disease; FTD, frontotemporal dementia; DS, Downs Syndrome; CJD, Creutzfeld-Jakob Disease. In all conditions analyzed, diseased brains showed hyperreactivity as well as hypertolerance.

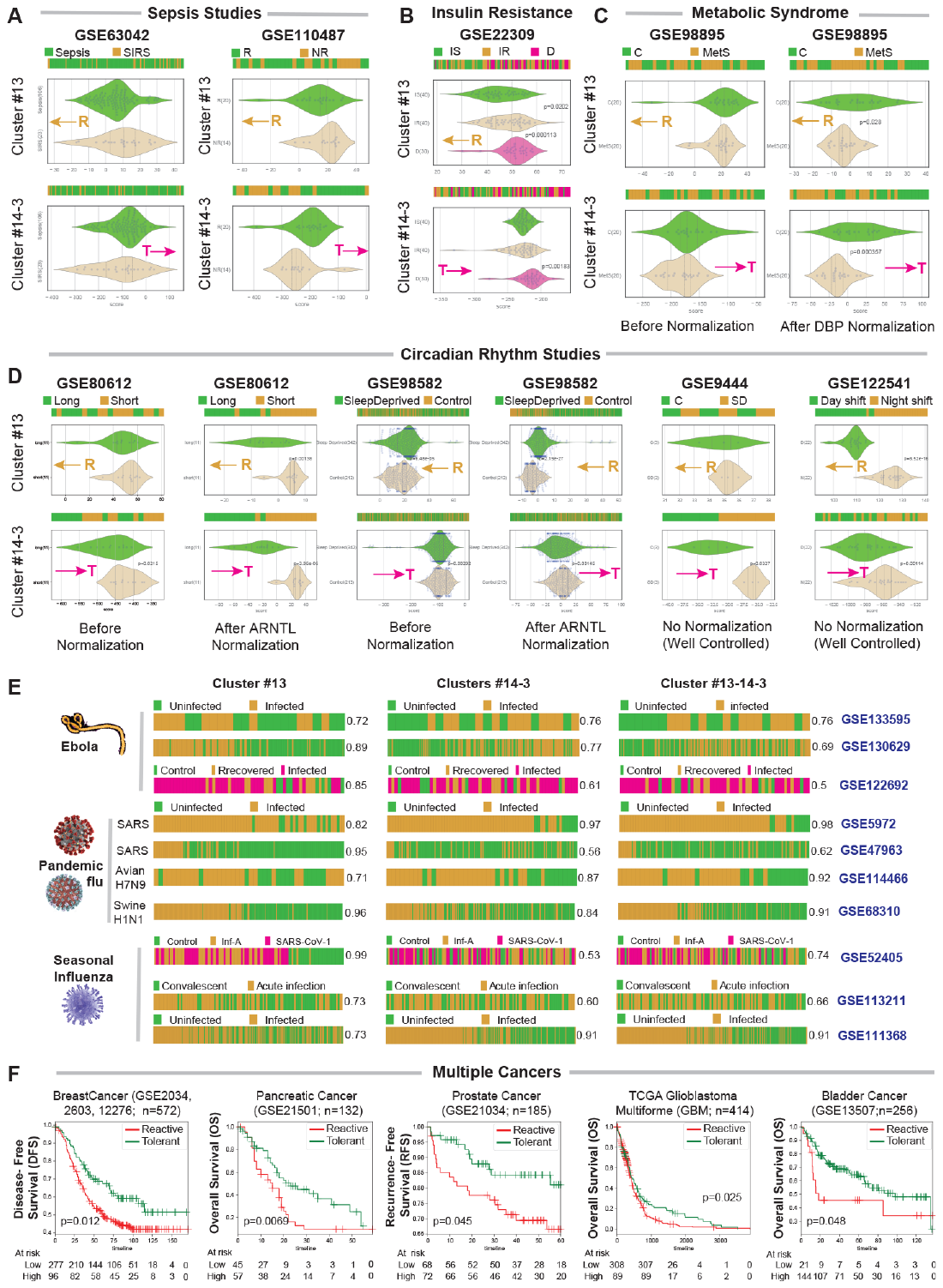

**Figure S7: Formal definitions of macrophage “reactivity” and “tolerance” identify pathologic states in sepsis, metabolic diseases and cancers.** **(A-D)**. Macrophage polarization states based on Cluster #13 and Boolean path #14-3 in sepsis (**A)** (*Left*) GSE63042; 106 sepsis [S] and 23 systemic inflammatory response syndrome [SIRS]; (*Right*) GSE110487; 20 responders [R] and 14 non-responders [NR]). (**B)** GSE22309; 40 insulin-sensitive [IS], 40 insulin-resistant [IR] and 30 Diabetic [D]). (**C**) GSE98895; 20 control [C] and 20 metabolic syndrome [MetS]) and in sleep disorders (**D**). **(C)** For metabolic syndrome, analysis is shown before and after normalization to DBP, a gene that controls circadian rhythm. **(D)** For sleep disorders study, two datasets (GSE80612, GSE98582) were normalized to ARNTL, a gene that controls circadian rhythm (analyses are shown before and after normalization) and the other two datasets (GSE9444, GSE122541) were analyzed without any normalization. **(E)** PBMC or lung tissue samples from humans infected with various respiratory viral infections and pandemics were classified based on either the levels of expression of genes in cluster #13 alone (*left*), clusters #14 and 3 (*middle*), or the path #13-14-3 (*right*). Numbers on the right side denotes ROC-AUC values for classification accuracy compared to uninfected controls. **(F)** Macrophage polarization based on Boolean path #13-14-3 is associated with outcome in several cancers, except in pancreatic cancer, where cluster #13 is prognostic.  P-values are derived from Welch's two-sided unpaired unequal variance Two Sample t-test between control and disease samples.

**Table S1: M0, M1 and M2 annotation in GSE134312**

M0:

| GSM300403 | GSM300400 | GSM249415 | GSM343826 | GSM300398 | GSM249395 |
| --- | --- | --- | --- | --- | --- |
| GSM249417 | GSM343820 | GSM249389 | GSM249371 | GSM343808 | GSM249375 |
| GSM249367 | GSM343802 | GSM343810 | GSM343828 | GSM249409 | GSM360182 |
| GSM249407 | GSM249381 | GSM360186 | GSM343814 | GSM300402 | GSM249403 |
| GSM249385 | GSM360139 | GSM343812 | GSM300405 | GSM343806 | GSM300406 |
| GSM213500 | GSM300404 | GSM249377 | GSM343816 | GSM343818 | GSM343804 |
| GSM360143 | GSM343824 | GSM343830 | GSM300401 | GSM343822 | GSM249399 |
| GSM249423 | GSM300399 | GSM115052 | GSM115053 | GSM115054 | |

M1:

| GSM300391 | GSM300393 | GSM300396 | GSM213511 | GSM300390 | GSM300389 |
| --- | --- | --- | --- | --- | --- |
| GSM300392 | GSM300394 | GSM300397 | GSM300395 | GSM115055 | GSM115056 |
| GSM115057 | |  |  |  |  |

M2:

| GSM183209 | GSM183196 | GSM183193 | GSM183165 | GSM183201 | GSM115058 |
| --- | --- | --- | --- | --- | --- |
| GSM115059 | GSM115060 | |  |  |  |

**Table S2: List of pattern recognition receptors on macrophages and their position within the Boolean network of macrophage processes.**

| **Receptors Family** | **Gene name** | **Ligands (Microbial/cells)** | **Microbes/binding partners and unique Function** | **Cluster Number** |
| --- | --- | --- | --- | --- |
| **TLR Family^1^** | TLR1 | Triacyl lipoprotein | Bacteria  Works in conjunction with TLR2 | Cluster 2 |
|  | TLR2 | Lipoproteins, Peptidoglycans,  Lipoteichoic acids | Bacteria, Virus | Cluster 2 and 5 |
|  | TLR3 | double stranded RNA | Virus | absent |
|  | TLR4 | LPS | Bacteria | Cluster 2 |
|  | TLR5 | Flagellin | Bacteria | Cluster 2, 3, 5 |
|  | TLR6 | Diacyl lipoprotein | Bacteria, Virus  Works in conjunction with TLR2 | absent |
|  | TLR7 | Single stranded RNA | Bacteria, Virus | Cluster 6 |
|  | TLR8 | Single stranded RNA | Bacteria, Virus | Cluster 2 |
|  | TLR9 | Unmethylated DNA with the CpG motif | Bacteria, Virus, protozoa | absent |
|  | TLR10 | Unknown | Unknown | absent |
|  | TLR11 | Profilin like molecule | Protozoa | absent |
|  | TLR12 | Profilin | Protozoa | absent |
|  | TLR13 | 23s ribosomal RNA | Bacteria | absent |
| **C-type Lectin Receptor (CLR) family^2^** | Type 1-DEC205/CD205 | Binds apoptotic and necrotic cells |  | absent |
|  | Type 1-Macrophage mannose receptor (MMR)/MRC1 | Mannose, Fucose, N-acetyl glucosamine or glucose | binds mannose on the surface of pathogenic viruses, bacteria, and fungi so that they can be neutralized by phagocytic engulfment. | Cluster2 |
|  | Type 2-Dectin 1/CLEC7A | Binds beta glucan of fungal cell wall | Phagocytose live yeast and zymosan from fungi | Cluster 2 |
|  | Type 2-Dectin 2/CLEC6A | Binding with Pneumocystis major surface glycoprotein/ glycoprotein A (Msg/gpA) |  | absent |
|  | Type 2-Mincle/CLEC4E | α-mannose, Cord factor | FcRγ-coupled CLR that bind to mycobacterial cord factor as well as certain fungal species | absent |
|  | Type 2-DC-Sign/CD209 | Binds mycobacteria- 4 known ligands-DnaK, 60 kDa chaperonin-1 (Cpn60.1), GAPDH and lipoprotein lprG. (PMID: 21203928) |  | Cluster 3 |
|  | Type 2-DNGR1/CLEC9A | Lectin |  |  |
|  | MBL | Array of carbohydrates |  |  |
|  | MGL | N-acetylgalactosamine  (GalNAc) residues |  |  |
| **Scavenger Receptor^3^** | SRA  (SCARA1-5) | Lipid A of LPS; soluble LTA from Streptococcus pyogenes | Phagocytic receptor helps in the non-opsonic uptake of bacteria | SCARA3 in cluster 1 |
|  | MARCO | Soluble LPS and LTA, CpG DNA, intact Gram-positive and Gram-  -negative bacteria | Host defense against S pneumonae infection in mouse model | absent |
|  | CD36 | Bacterial membrane, LPS |  | absent |
|  | SRB1/SCARB1 | Binds high density lipoprotein |  | Cluster 1 and 3 |
| **Nod like receptor (NLR)^4^** | Nod1 | Di-amino-pimelic Acid | Cytosolic sensor, activates host signaling by Rip2 | Cluster 1 |
|  | Nod2 | Muramyl dipeptide |  | Cluster 2 and 5 |
|  | Naip | Flagellin |  | absent |
|  | NLRC3 | Unknown |  | absent |
|  | NLRC4 | Flagellin |  | absent |
|  | NLRP1 | Muramyl dipeptide |  | absent |
| **Receptor for advanced glycation end-products (RAGE)^5^** | RAGE | Multi ligands: AGEs, HMGB1, S100s, beta sheet fibrils, DNA, RNA |  | Cluster 1 |
| **Apoptotic cell recognition by direct binding^6^** | BAI1 | Phosphatidyl Serine, bacterial LPS | GPCR- ELMO1, DOCK, Rac1 | Cluster 1 |
|  | TIM1 | Phosphatidyl Serine, hepatitis virus A | Signal via Fyn Kinase |  |
|  | TIM4 | Phosphatidyl Serine | Indirect via integrins | Absent |
|  | Stabilin 2 | Phosphatidyl Serine | ITIM | Cluster 1 |
|  | CD300 | Phosphatidyl Serine | Via GULP | Absent |
| **Apoptotic cell recognition by indirect binding^7^** | MER- type I receptor tyrosine kinase | Protein S (PROS1) |  | Cluster 1 |
|  | AXL- type I receptor tyrosine kinase | GAS6 |  | Absent |
|  | Tyro3- type I receptor tyrosine kinase | GAS6, PROS1 |  | Cluster 1 |
|  | SCARF | C1q |  | absent |
|  | Integrin aMb2 | C1q |  | Cluster 1 |
|  | Integrin avb3 | MFG-E8 | CRKII, DOCK180, Rac1 | Cluster 2 |
|  | Integrin avb5 | MFG-E8 | FAK, DOCK180, Rac1 | Cluster 2 |
|  | CD36 | oxidized low-density lipoprotein (oxLDL), anionic phospholipids, long-chain fatty acids, thrombospondin-1 (TSP1), fibrillar β-amyloid | Fyn, PYK2 | absent |

1-7 collected from Fox & Das, Book Chapter, 2015

^1^ PMID: 23284045; PMID: 12524386, PMID: 11905821; PMID: 20303872

^2^ PMID: 29497419; PMID: 32060644

^3^ PMID: 30318962

^4^ PMID: 20303872

^5^ PMID: 15488742

^6,7^ PMID: 26683144; PMID: 29400998

**Table S3: Datasets identified in NCBI GEO database in which the expression or function of targets in cluster #13 or #14 were experimentally manipulated in cells of the myeloid lineage.**

| Genes | GSEID | Model/approach | Species | Predicted outcome | Observed outcome |
| --- | --- | --- | --- | --- | --- |
| AIM2 | GSE24155 | Overexpression, HCT116 (colorectal tumour cell line) | Human | Hyper-reactivity | Hyper-reactive |
| Oas2 | GSE69397 | KO mice | Mouse | Hyper-reactivity | Hyper-reactive |
| IL15/IL15R | GSE132270 | Recombinant IL15 Monocytes | Human | Hyper-reactivity | Hyper-reactive |
| IRF1 | GSE66461 | KO mice | Mouse | Hyper-tolerence | Hyper-tolerant |
| IFIT2 | GSE33678 | KO mice | Mouse | Hyper-tolerence | Hyper-tolerant |
| STAT3 | GSE131300 | Inhibitor ruxolitinib, Primary human macrophage | Human | Hyper-tolerence | Hyper-tolerant |
| STAT3 | GSE131300 | KO mice, BMDM | Mouse | Hyper-tolerence | Hyper-tolerant |
| Oas2 (I405N) | GSE69397 | KO mice | Mouse | Hyper-tolerence | Hyper-tolerant |
| USP18 | [GSE61501](https://www.ncbi.nlm.nih.gov/geo/query/acc.cgi?acc=GSE61501) | KO mice, Microglia | Mouse | Hyper-tolerence | Hyper-tolerant |
| CCDC88A |  | KO mice, Peritoneal macrophage | Mouse | Hyper-reactivity | Hyper-reactive |
| TSC2 | [GSE77075](https://www.ncbi.nlm.nih.gov/geo/query/acc.cgi?acc=GSE77075) | KO mice, BMDM | Mouse | Hyper-reactivity | Hyper-reactive |
| RNF5 | [GSE127753](https://www.ncbi.nlm.nih.gov/geo/query/acc.cgi?acc=GSE127753) | KO mice | Mouse | Hyper-reactivity | Hyper-reactive |
| MLL | GSE129637 | NPM1c knock-in cells, inhibitor VTP-50469 | Mouse | Hyper-reactivity | Hyper-reactive |
| MLL | GSE129637 | AML cell lines carrying NPM1c mutations, inhibitor VTP-50469 | Human | Hyper-reactivity | Hyper-reactive |
| MLL | GSE129637 | inhibitor VTP-50469 |  | Hyper-reactivity | Hyper-reactive |
| EZH1 | [GSE110925](https://www.ncbi.nlm.nih.gov/geo/query/acc.cgi?acc=GSE110925) | KO mice | Mouse | Hyper-reactivity | Hyper-reactive |
| NFX1, PCBP2, EEF2, HNRNPA1 | ENCODE | ShRNA | Human | Hyper-reactivity | Hyper-reactive |
